# Brain Capillary Pericytes are Metabolic Sentinels that Control Blood Flow through K_ATP_ Channel Activity

**DOI:** 10.1101/2022.03.14.484304

**Authors:** Ashwini Hariharan, Colin D. Robertson, Daniela C.G. Garcia, Thomas A. Longden

**Affiliations:** Department of Physiology, School of Medicine, University of Maryland, Baltimore, MD, USA; Department of Pharmacology, School of Medicine, University of Maryland, Baltimore, MD, USA

**Keywords:** Pericytes, endothelial cells, capillaries, neurovascular coupling, functional hyperemia, K_ATP_ channels, K_IR_ channels, cerebral blood flow, glucose, energy, metabolism

## Abstract

Capillary pericytes and their processes cover ∼90% of the total length of the brains capillary bed. Despite their abundance, little is known of pericyte function, and their contributions to the control of brain hemodynamics remain unclear. Here, we report that deep capillary pericytes possess a mechanistic ‘energy switch’ that, when activated by a decrease in glucose, elicits robust K_ATP_ channel activation to increase blood flow and protect energy substrate availability. We demonstrate that pharmacological activation of K_ATP_ channels profoundly hyperpolarizes capillary pericytes and leads to dilation of upstream penetrating arterioles and arteriole-proximate capillaries covered with contractile pericytes, leading to an increase in local capillary blood flow. Stimulation of a single capillary pericyte with a K_ATP_ channel agonist is sufficient to evoke this response, which is mediated via K_IR_ channel-dependent retrograde propagation of hyperpolarizing electrical signals. Genetic inactivation of pericyte K_ATP_ channels via expression of a dominant-negative version of K_IR_6.1 eliminates these effects. Critically, we show that lowering extracellular glucose below 1 mM evokes dramatic K_ATP_ channel-mediated pericyte hyperpolarization. Inhibiting glucose uptake by blocking GLUT1 transporters *in vivo* also activates this energy switch to increase pericyte K_ATP_ channel activity, dilate arterioles and increase blood flow. Together, our findings recast capillary pericytes as metabolic sentinels that respond to local energy deficits by robustly increasing blood flow to protect metabolic substrate delivery to neurons and prevent energetic shortfalls.

## INTRODUCTION

Blood flow provides the oxygen and glucose that are critical for the metabolic processes that underpin brain function and health. Thus, precise control of cerebral hemodynamics is essential to meet the moment-to-moment energetic needs of neurons and glia. The brains vascular system is fed by pial arteries, which originate at the circle of Willis and course over the surface of the brain before branching orthogonally to give way to penetrating arterioles (PAs) that dive into the parenchyma^1^. PAs, in turn, branch to give rise to a tortuous capillary network that is covered by a diverse population of pericytes. At the first point of the PA-to-capillary transition, mural cells termed ‘pre-capillary sphincters’ are found which exert dynamic control of blood flow into the capillary bed by virtue of their α-smooth muscle actin (SMA) expression^2^. Adjacent to this, the initial 3-4 branches of the capillary network are collectively referred to as a ‘transitional segment’^3^ due to their coverage by contractile pericytes that express α-SMA and these cells are capable of rapidly regulating the diameter and therefore blood flow of the underlying vessel^4–6^. Immediately downstream of the α-SMA terminus are mesh pericytes, and deeper in the capillary bed (from approximately the 5^th^ branch and above), the abluminal surface of the capillaries is adorned by the processes and cell bodies of thin-strand pericytes^4,7^. The latter extend long, narrow processes which stretch in some cases for hundreds of microns along the walls of local capillaries, coming into close apposition with the arborizations of neighboring thin-strand pericytes^8^. They also form ‘peg-socket’ junctions with the underlying ECs, which are thought to be the sites of gap junction coupling, permitting the ready exchange of molecules and charge between these cells^9–11^.

Pericytes contribute to multiple physiological processes including regulation of blood brain barrier permeability and modulation of endothelial cell (EC) gene expression^12,13^. As they are ideally positioned to mediate communication between the blood and brain parenchyma, it has been suggested that these cells play a critical role in control of hemodynamics. A growing body of evidence indicates that contractile pericytes of the transitional segment play a key role in rapidly regulating the diameter of the underlying capillaries^2,5,14–18^. However, the mechanisms for blood flow control by thin-strand pericytes have not been defined. Emerging evidence suggests that subtle contractile processes in these cells may regulate capillary diameter and therefore local capillary blood flow^6^. Here, we show that the electrical activity of thin-strand pericytes alone is sufficient for robust, remote blood flow control by these cells via communication with the underlying endothelium. We recently surveyed the molecular expression of ion channels and G protein-coupled receptors (GPCRs) in thin-strand pericytes of the brain^19^. The vascular form of the ATP-sensitive potassium (K^+^; K_ATP_) channel, composed of inward rectifier K^+^ (K_IR_) 6.1 and sulfonylurea receptor (SUR) 2 subunits is the most highly expressed ion channel subtype in these cells and accounts for almost half of their relative expression of all ion channel genes^20,21^. As K_ATP_ channels are found in a range of tissues where they play a major role in coupling metabolism to membrane electrical activity^22,23^, we hypothesized that they may play a similar role in brain pericytes, linking local metabolic substrate availability to membrane hyperpolarization and, ultimately, blood flow control.

We demonstrate here that deep capillary pericytes control local blood flow via K_ATP_ channel-mediated electrical signaling. Our results indicate that pericyte K_ATP_ channels are the molecular cornerstone of an ‘energy switch’ mechanism, wherein a fall in glucose availability below a key threshold evokes a K_ATP_ channel-mediated blood flow increase to replenish energy substrate delivery to neurons and glia. Our data thus recast thin-strand pericytes as metabolic sentinels that dynamically modulate blood flow to ensure that the energy substrates required to support ongoing neuronal function are continually provided.

## RESULTS

### K_ATP_ channel activation increases arteriolar diameter and capillary blood flow *in vivo*

To examine the role of K_ATP_ channels in the control of brain blood flow, we began by visualizing the vascular network of a volume of cortex through a cranial window preparation in mice anesthetized with urethane and alpha-chloralose (**Fig. 1A**). We identified pial arteries on the brains surface and their arising PAs branching perpendicularly into the tissue by their morphology in relation to nearby veins, and imaged these PAs and their daughter capillaries down to at least the 5^th^ branch of the capillary bed (**Fig. 1A**). *In vivo*, these arteries constrict partially in response to intravascular pressure^24^, establishing a baseline of myogenic tone from which diameter can be bidirectionally modulated to adjust blood flow. Under our conditions, we found that PAs had 46.36 ± 2.84% tone (*n* = 8 arterioles from 5 mice), calculated by comparing baseline arteriolar diameter to passive diameter in the absence of extracellular calcium (Ca^2+^) and the presence of the voltage-dependent Ca^2+^ channel blocker diltiazem (200 μM) (**Supplementary Fig. 1A**). We also measured the tone of the 1^st^ to 4^th^ order branches of the capillaries of the transitional segment under the same conditions, covered with the cell bodies and processes of contractile pericytes. These branches collectively averaged 39.35 ± 2.18 % tone at baseline, which was not different to the tone of PAs and did not differ by branch order (*n* = 35 capillaries from 5 mice, **Supplementary Fig. 1A**). To examine the influence of K_ATP_ channels on capillary and arteriole diameter and blood flow, we assessed the effects of pharmacologically modulating these channels through superfusion of agents over the cranial surface. Strikingly, we found that application of 10 μM pinacidil, a selective K_ATP_ channel opener, produced near-maximal dilation of both PAs and transitional segment capillaries (**Fig. 1B-H**), indicating that K_ATP_ channels can exert a strong influence on the vasculature. In turn, these substantial increases in diameter translated into profound elevation of capillary blood flow, measured as red blood cell (RBC) flux using a high-frequency line scanning approach (**Fig. 1I, J**). To determine whether K_ATP_ channel activity contributes to basal blood flow, we applied the K_ATP_ channel blocker glibenclamide (10 μM) to the cranial surface. We observed no change in the diameter of the PA or 1^st^ – 4^th^ order capillaries to this maneuver, and no change in capillary blood flow (**Supplementary Fig. 2**), suggesting that vascular K_ATP_ channel activity in the brain is minimal under our resting conditions.

**Figure 1.**
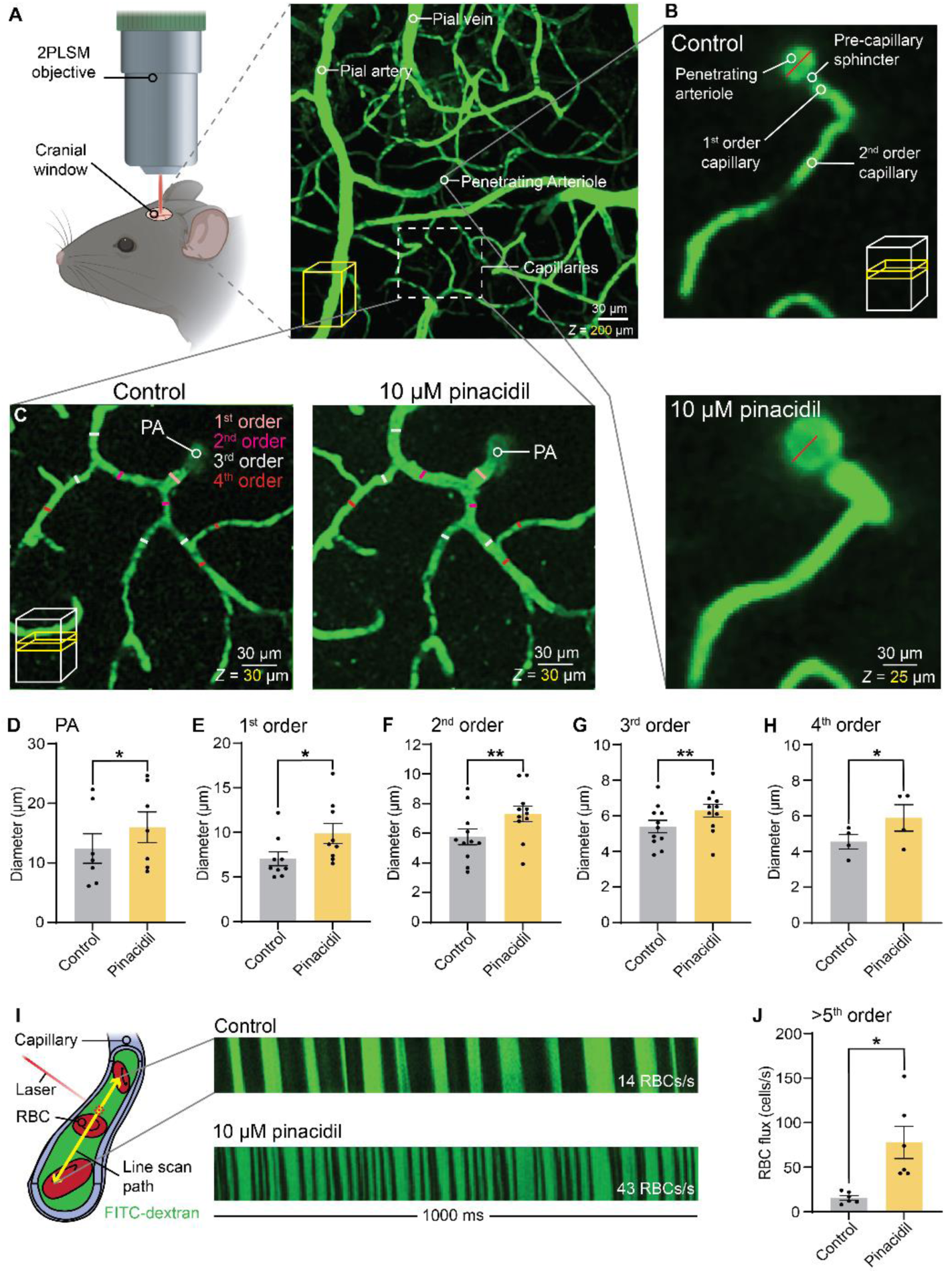
K_ATP_ channel activation increases arteriole diameter and capillary blood flow *in vivo*. **(A)** *In vivo* imaging set-up. *Left:* A cranial window was made over the somatosensory cortex and imaged using two-photon laser scanning microscopy. *Right:* Imaging field of the cortical vasculature containing FITC-dextran showing a pial vein and artery, a penetrating parenchymal arteriole (PA) and downstream capillaries. **(B)** A PA with a pre-capillary sphincter and its downstream 1^st^ and 2^nd^ order capillaries. *Top:* Baseline diameter of the PA (red line). *Bottom:* Dilation of the PA after application of 10 μM pinacidil (red line: baseline diameter). The capillaries in view also dilated to this maneuver. **(C)** A PA and its downstream 1^st^-4^th^ order capillaries. *Left:* Baseline diameters of 1^st^-4^th^ order capillaries indicated by respective colored lines. *Right:* The same 1^st^-4^th^ order capillaries, which dilated after application of 10 μM pinacidil. **(D-H)** Summary data, analyzed using paired Student’s *t*-test, showing application of 10 μM pinacidil produced a significant dilation of the **(D)** PA (n = 7 vessels, 7 mice, \**P* = 0.01, t_6_ = 3.697), **(E)** 1^st^ order capillary (n = 9 vessels, 7 mice, \**P* = 0.017, t_8_ = 2.987), **(F)** 2^nd^ order capillary (n = 11 vessels, 6 mice, \*\**P* = 0.0029, t_10_ = 3.921), **(G)** 3^rd^ order capillary (n = 11 vessels, 6 mice, \*\**P* = 0.0011, t_10_ = 4.513) and **(H)** 4^th^ order capillary (n = 4 vessels, 4 mice, \**P* = 0.0326, t_3_ = 3.772). **(I)** Line scanning strategy used to measure blood flow in higher order capillaries. *Top-right:* Kymograph taken at baseline displaying RBCs passing through the line-scanned capillary as dark shadows against the green fluorescence of FITC-containing plasma. *Bottom-right:* Kymograph of the same capillary post-pinacidil application showing a dramatic increase in RBC flux. **(J)** Summary of RBC flux responses showing significant hyperemia to 10 μM pinacidil (n = 6 vessels, 3 mice, \**P* = 0.0188, t_5_ = 3.424, paired Student’s *t*-test).

### Pericytes transmit K_ATP_ channel-mediated electrical signals via the endothelium to exert remote control over the diameter of upstream arterioles

The expression of K_ATP_ channels is relatively lower in SMCs and in arteriolar and capillary endothelial cells (ECs) of the brain compared to thin-strand pericytes^19–21,25^, and pharmacological maneuvers designed to activate these channels in isolated PAs do not lead to dilation^26^. Given that systems-level K_ATP_ channel activation evoked profound vasodilation and blood flow increases, we reasoned that the high expression of K_ATP_ channels in deep capillary thin-strand pericytes could be the primary source of hyperpolarizing signals that may then be relayed to upstream PAs to drive their vasodilation. To test this possibility, we maneuvered a pipette connected to a pressure-ejection system into the brain and positioned it next to a DsRed-positive thin-strand pericyte in *Cspg4*-DsRed mice (**Fig. 2A-C**). On average, targeted pericytes were 268.5 ± 25.9 μm from the upstream arteriole imaging site (n = 8 experiments, 8 mice). Consistent with our hypothesis, activation of K_ATP_ channels in a single pericyte by local pressure ejection of 10 μM pinacidil onto the pericyte cell body (**Fig. 2C**) evoked a rapid and substantial upstream arteriolar dilation (**Fig. 2D,E,G** and **Supplementary Movie 1**) which was accompanied by an increase in underlying capillary blood flow (**Fig. 2F,H**).

**Figure 2.**
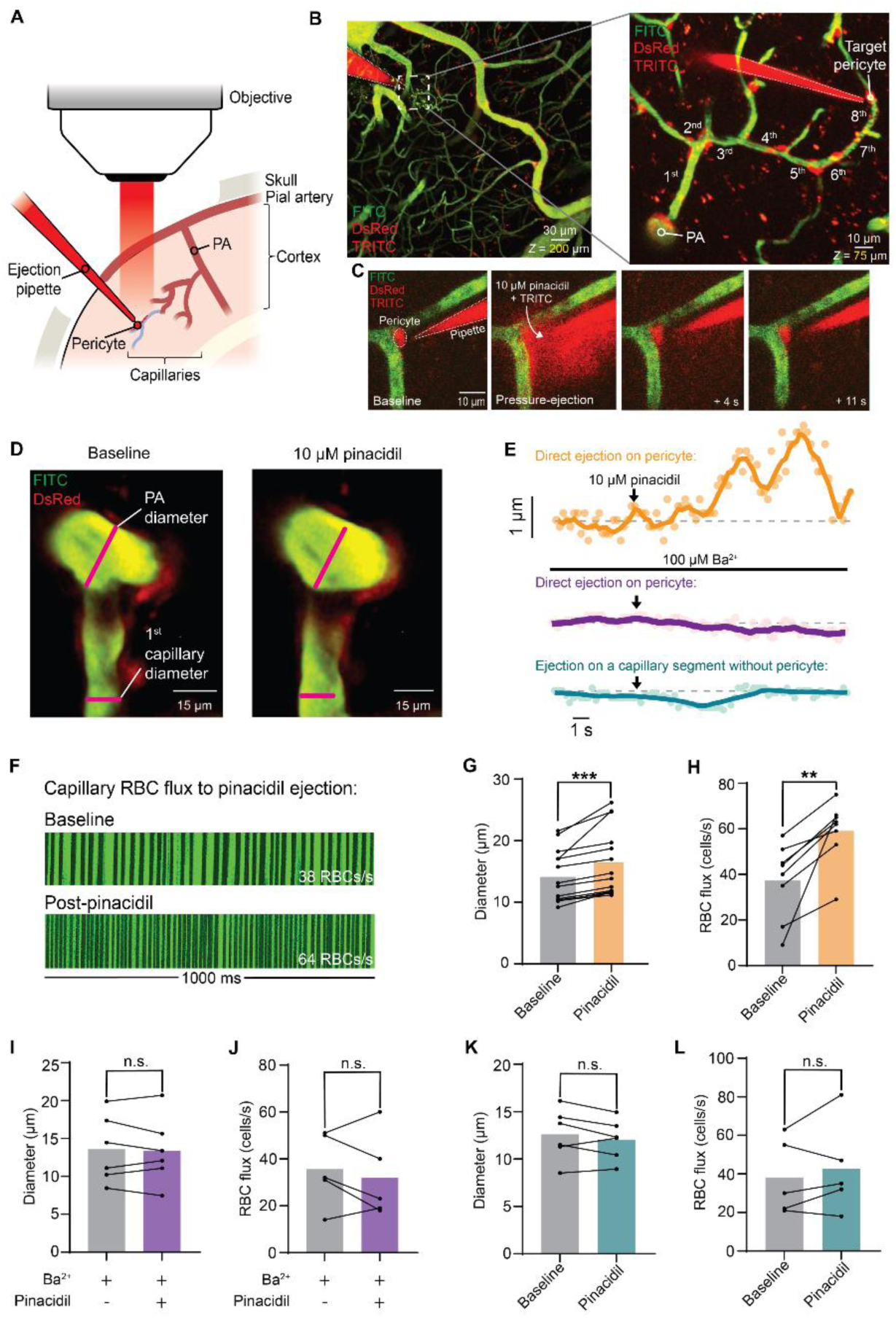
Capillary pericytes exert remote control over upstream PA diameter. **(A)** Cartoon illustrating the experimental strategy, showing an ejection pipette positioned next to a pericyte. **(B)** *Z*-projections of 3D volume acquisitions outlining the experimental strategy. *Left:* Vasculature containing FITC-dextran and a pipette with TRITC-dextran positioned within the cortex. *Right:* A PA and its downstream capillary network showing an ejection pipette containing TRITC-dextran with 10 μM pinacidil positioned next to a DsRed+ pericyte on an 8^th^ order capillary. **(C)** Depiction of the evolution (left to right) of TRITC diffusion (red) after pressure ejection of 10 μM pinacidil onto a DsRed+ pericyte. The brevity and low pressure of the ejection conditions (10 psi, 30 ms) ensured that the drug remained local. **(D)** Focal stimulation of capillary pericytes with 10 μM pinacidil dilates the connected upstream PA. *Left:* PA and 1^st^ order capillary diameter at baseline indicated by magenta lines. *Right:* Peak dilation of the same PA and 1^st^ order capillary after pinacidil-ejection on the downstream pericyte. **(E)** Representative time courses showing PA dilation to direct stimulation of a pericyte with pinacidil (orange, *top*), but no change in PA diameter when pinacidil was applied in the presence of the K_IR_ channel blocker Ba^2+^ (purple, *middle*) or when pinacidil was ejected onto a segment of capillary without a pericyte cell body (blue, *bottom*). **(F)** 1-s kymograph segments showing raw RBC flux of a >5^th^ order capillary at baseline, and hyperemia after pinacidil was ejected onto the overlying pericyte. **(G)** Summary of PA diameter changes after pinacidil-ejection on a downstream pericyte (n = 14 paired measurements, 13 mice, \*\*\**P* = 0.0007, t_13_ = 4.439, paired Student’s *t*-test). **(H)** Summary capillary RBC flux responses to pinacidil applied directly to a pericyte (n = 8 paired measurements, 4 mice, \*\**P* = 0.0045, t_7_ = 4.108, paired Student’s *t*-test). **(I)** Summary data showing PA diameter after pinacidil-stimulation of a pericyte in the presence of Ba^2+^ (n = 6 paired measurements, 6 mice, *P* = 0.6981, t_5_ = 0.411, paired Student’s *t*-test). **(J)** Summary blood flow data showing RBC flux before and after pinacidil-stimulation of a pericyte in the presence of Ba^2+^ (n = 5 paired measurements, 5 mice, *P* = 0.4613, t_4_ = 0.814, paired Student’s *t*-test). **(K)** Summary data showing PA diameter changes on stimulation of a capillary segment without a pericyte cell body with pinacidil (n = 6 paired measurements, 6 mice, *P* = 0.2162, t_5_ = 1.415, paired Student’s *t*-test). **(L)** Summary of RBC flux responses before and after stimulation of a capillary segment without pericytes with pinacidil (n = 5 paired measurements, 5 mice, *P* = 0.394, t_4_ = 0.9543, paired Student’s *t*-test).

Pericytes are intricately associated with adjacent ECs via peg-socket processes which are thought to be the sites of gap junction coupling between these two cell types^11,19,27^. Accordingly, we reasoned that signals originating in pericytes may be transmitted upstream via connected underlying ECs. We previously identified an EC-mediated regenerative electrical signaling mechanism dependent on inward-rectifier K^+^ (K_IR_2.1) channels that transmits dilatory signals from deep within the capillary bed to upstream PAs^28^. Interestingly, blocking K_IR_2.1 channels by the application of 100 μM barium (Ba^2+^) to the cortical surface prior to pinacidil ejection on a pericyte abolished this increase in arteriolar diameter and capillary blood flow (**Fig. 2E,I,J**), suggesting that K_ATP_ channel-initiated hyperpolarization modulates electrical signaling through the capillary bed to produce its effects. Consistent with pericytes being the locus of pinacidil-evoked vasodilatory drive, the ejection of this agent onto a segment of capillary lacking a pericyte soma had no effect on arteriolar diameter or local blood flow (**Fig. 2 E,K,L**). As expected, diameter and blood flow were also unchanged when pericytes were stimulated with vehicle (artificial cerebrospinal fluid (aCSF) containing 0.3 mg/mL TRITC-dextran) (**Supplementary Fig. 3**). Importantly, direct stimulation of the PA with 10 μM pinacidil also had no effect on diameter (**Supplementary Fig. 4**), which aligns with previous observations of a lack of response of these arterioles to K_ATP_ agonists^26^ and buttresses the conclusion that pericyte K_ATP_ channels exert remote control of upstream PA diameter.

### Expression of a dominant-negative mutant of the vascular K_ATP_ channel eliminates pericyte-mediated dilations and hyperemia

To unequivocally confirm the central role of pericyte K_ATP_ channels in control of blood flow and upstream PA diameter to pinacidil, we deployed mice that express a dominant-negative form of the K_IR_6.1 subunit in which a Gly-Phe-Gly motif of the K^+^ selectivity filter is mutated to a non-functional alanine triplet (K_IR_6.1^AAA^), which in turn eliminates K_ATP_ currents^29,30^. Expression of K_IR_6.1^AAA^ was controlled by tamoxifen-inducible Cre-recombinase under the *Cspg4* promoter to selectively suppress K_ATP_ channel activity in pericytes and SMCs. In this line, a floxed region containing the sequence for enhanced green fluorescent protein (eGFP) upstream of a stop codon is expressed under basal conditions, precluding expression of the downstream K_IR_6.1^AAA^ sequence without Cre-recombinase activity. When recombination is induced, eGFP along with the stop codon are excised, permitting K_IR_6.1^AAA^ expression (**Fig. 3A**). Accordingly, induction of Cre activity in *Cspg4*-Cre-K_IR_6.1^AAA^ mice by 4-hydroxy tamoxifen (4-OHT) eliminated eGFP expression in capillary pericytes, while eGFP expression was retained in adjacent ECs (**Fig. 3B,C**), indicating successful cell type-selective expression of the K_IR_6.1^AAA^ construct. To then reveal pericytes with inactive K_ATP_ channels, we applied NeuroTrace 500/525 (NT500/525)^31^, to the cranial surface which specifically stained thin-strand pericytes (**Fig. 3C**). Pressure-ejecting pinacidil onto thus identified eGFP-negative, NT500/525-positive pericytes did not produce an increase in upstream PA dilation or local capillary blood flow in *Cspg4*-Cre-K_IR_6.1^AAA^ mice (**Fig. 3C,D,I,J**), indicating that functional K_ATP_ channels in pericytes are essential for these responses. However, Cre control (K_IR_6.1^AAA^ mice given 4-OHT) and vehicle control (*Cspg4*-Cre-K_IR_6.1^AAA^ mice given a 90:10% mixture of corn oil:ethanol) groups still demonstrated significant PA dilation (11-13%) and capillary RBC flux still increased (32-38%) to these maneuvers (**Fig. 3D,E-H**). Thus, pericytes are the primary site of K_ATP_-mediated upstream arteriolar dilation and local capillary hyperemia *in vivo*.

**Figure 3.**
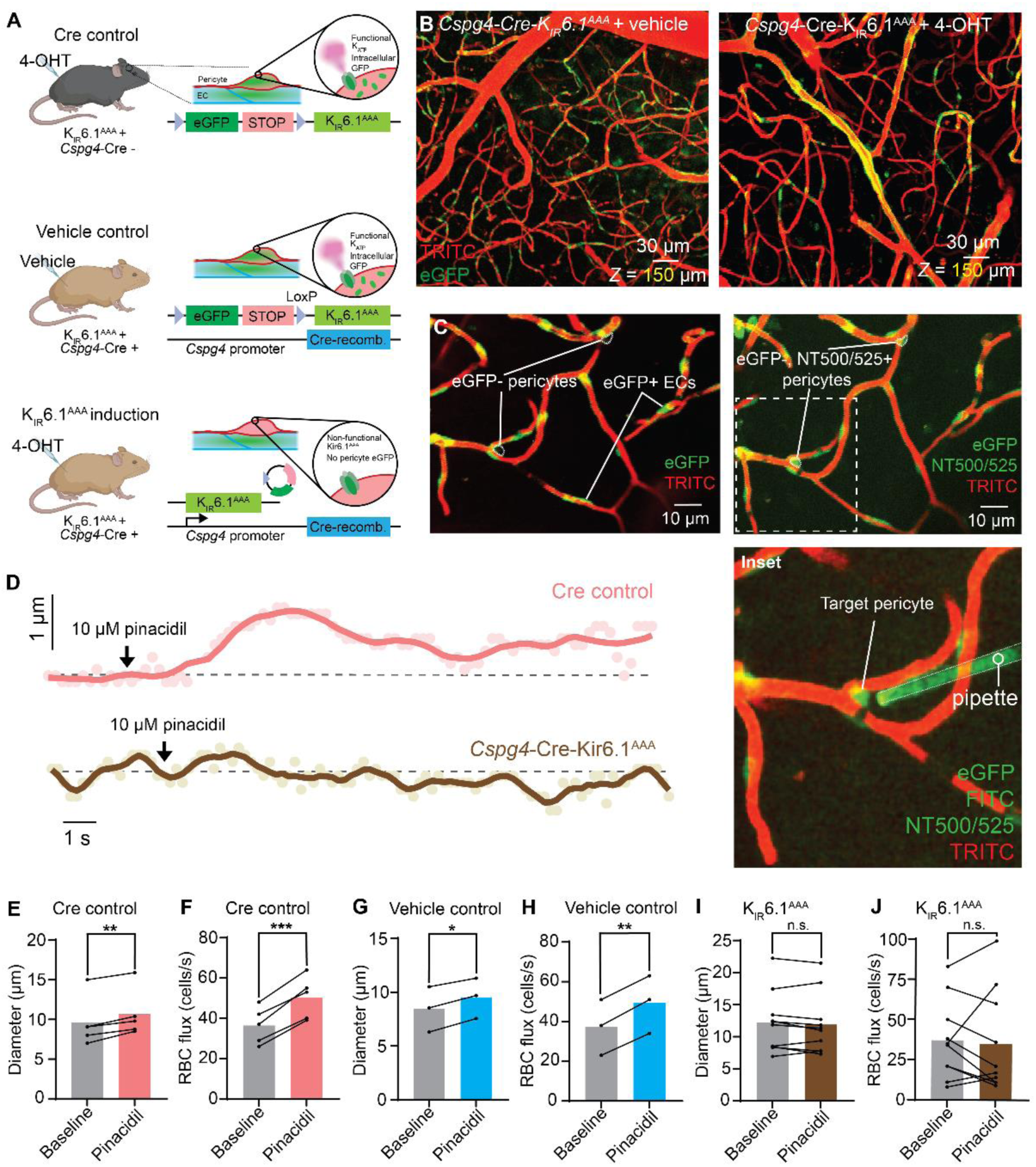
Capillary pericytes are the locus of K_ATP_ channel-mediated control of blood flow. **(A)** Pericyte K_ATP_ channels were genetically inactivated by crossing mice possessing a modified K_IR_6.1 subunit (K_IR_6.1^AAA^) with *Cspg4*-Cre mice. Cre control mice (K_IR_6.1^AAA^+, Cre -) and K_IR_6.1^AAA^+ mice (K_IR_6.1^AAA^+, Cre +) were given 4-hydroxytamoxifen (4-OHT), whereas vehicle control mice (K_IR_6.1^AAA^+, Cre +) were given vehicle. **(B)** Successful inactivation of the K_IR_6.1 subunit was evidenced by elimination of eGFP signal in pericytes. *Left:* Representative Z-projection from a vehicle control mouse. *Right:* Representative Z-projection from a tamoxifen-induced *Cspg4*-Cre-K_IR_6.1^AAA^ mouse showing fewer eGFP+ cells. **(C)** Experimental strategy to identify and target inactivated K_ATP_ channels in pericytes. *Left:* A *Cspg4*-Cre-K_IR_6.1^AAA^ mouse, with eGFP+ endothelial cells, and pericytes lacking eGFP signal, indicating successful K_IR_6.1^AAA^ induction. *Right:* The location of eGFP-negative pericytes was determined using the *in vivo* pericyte-specific dye Neurotrace (NT) 500/525. *Inset:* A pipette containing FITC and 10 μM pinacidil positioned next to an eGFP-, NT 500/525+ pericyte. **(D)** Example traces of PA diameter showing dilation to downstream ejection of pinacidil onto a capillary pericyte in a Cre-control mouse (pink) and a lack of response in *Cspg4*-Cre-K_IR_6.1^AAA^ mice (brown). **(E-J)** Summary data of changes in PA diameter and blood flow to focal application of pinacidil onto a capillary pericyte across different experimental groups. **(E)** PA diameter changes in Cre-control mice (n = 5 paired measurements, 5 mice, \*\**P* = 0.0026, t_4_ = 6.684). **(F)** RBC flux changes in Cre-control mice (n = 5 paired measurements, 5 mice, \*\*\**P* = 0.0006, t_4_ = 9.908). **(G)** PA diameter changes in vehicle control mice (n = 3 paired measurements, 3 mice, \**P* = 0.0157, t_2_ = 7.883). **(H)** RBC flux changes in vehicle control mice (n = 3 paired measurements, 3 mice, \*\**P* = 0.0023, t_2_ = 20.78). **(I)** PA diameter changes in *Cspg4*-Cre-K_IR_6.1^AAA^ mice (n = 10 paired measurements, 10 mice, *P* = 0.3054, t_9_ = 1.087). **(J)** RBC flux changes in *Cspg4*-Cre-K_IR_6.1^AAA^ mice (n = 10 paired measurements, 10 mice, *P* = 0.7249, t_9_ = 0.3631). All data were analyzed using paired Student’s *t*-test.

### An energy-sensing switch couples decreases in local energy substrate availability to membrane hyperpolarization via K_ATP_ channel activity

Having established that pericyte K_ATP_ channels can exert a profound influence over PA diameter and local blood flow, we next turned our attention to the mechanisms through which K_ATP_ channels may be engaged. In other tissues, K_ATP_ channels play a critical role in coupling metabolism to membrane electrical activity, and are sensitive to the local level of glucose^32^. We thus hypothesized that pericytes might sense fluctuations in glucose levels in the brain and respond to decreases in glucose availability with K_ATP_ channel-mediated electrical signals.

Glucose concentration in bulk cerebrospinal fluid is ∼4 mM^33^, whereas parenchymal glucose has been measured in the range of 0.25-2.5 mM across a range of studies^34–42^. Accordingly, we wondered whether subtle changes in local glucose concentration in this range would influence the degree of K_ATP_ channel activity and thus modulate pericyte membrane potential (Vm). To explore the relationship between glucose and pericyte Vm, we applied a series of decreasing glucose concentrations to isolated capillaries from *Cspg4*-DsRed mice with intact thin-strand pericytes, and measured Vm using microelectrode impalements (**Fig. 4A**). Across all conditions of replete glucose (4 mM), pericyte Vm averaged -36 mV (22 cells, 10 mice; **Fig. 4B,E,F,I**).

**Figure 4.**
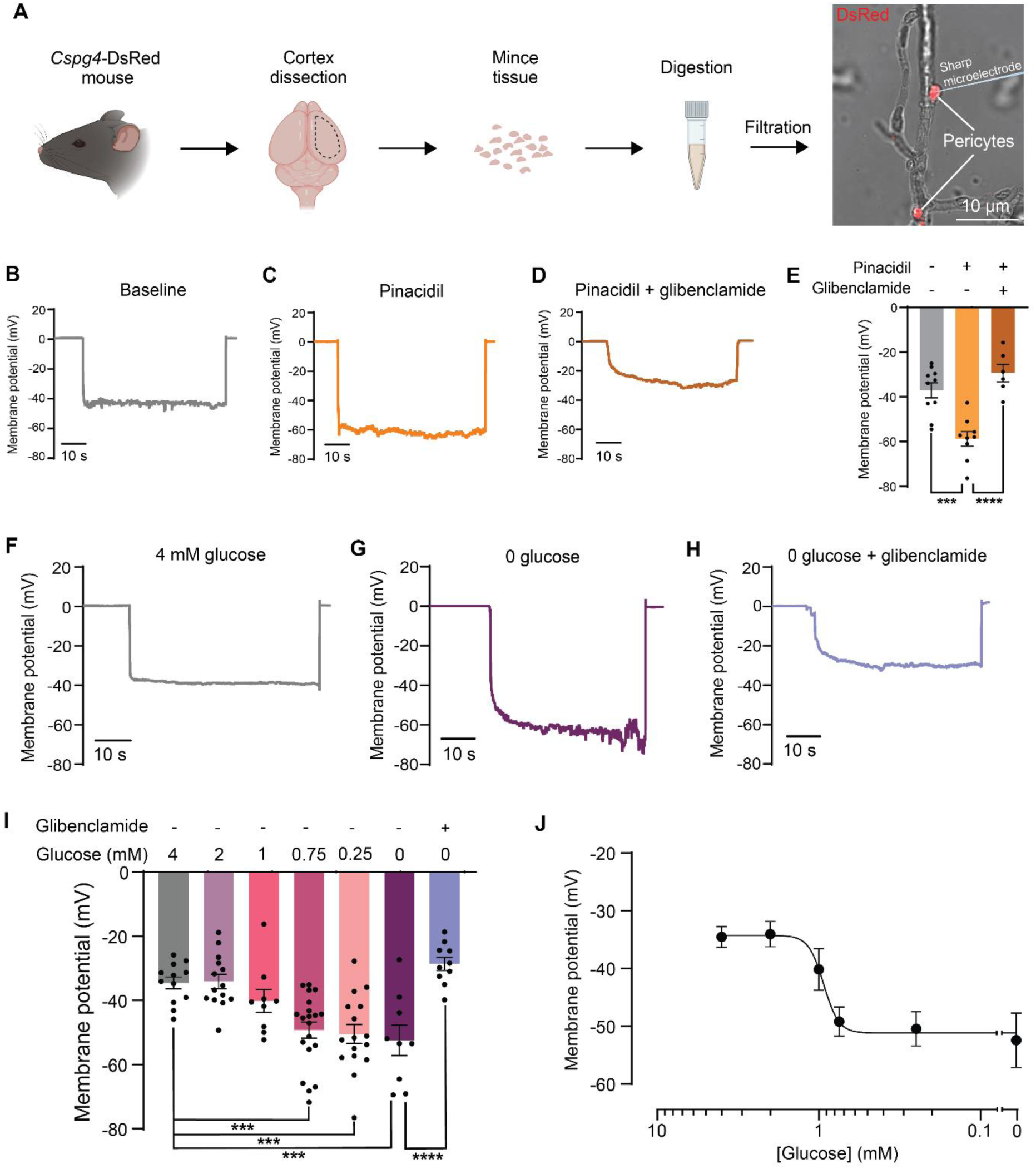
Lowering glucose activates K_ATP_ channels to hyperpolarize pericytes. **(A)** Overview of cell isolation and impalement. *Left to right:* Pericytes were isolated by dissecting and mincing cortical tissue from a *Cspg4*-DsRed mouse. Minced pieces were sequentially digested, homogenized and filtered to yield capillary fragments with DsRed-positive pericytes. **(B-D)** Example traces of Vm measurements at baseline **(B)**, with 10 μM pinacidil **(C)**, and with 10 μM pinacidil in the presence of 10 μM glibenclamide **(D). (E)** Summary data showing pinacidil hyperpolarizes pericyte Vm, and glibenclamide blocks this effect (baseline (10 cells, 5 mice) vs. pinacidil (9 cells, 4 mice): \*\*\**P* = 0.002, t_49_ = 4.453; pinacidil vs. pinacidil + glibenclamide (6 cells, 4 mice): \*\*\*\**P* < 0.0001, t_49_ = 5.278, One-way ANOVA with Sidak’s multiple comparison test). **(F-H)** Example traces of Vm measurements with 4 mM bath glucose **(F)**, with 0 bath glucose **(G)** and under 0 glucose conditions with the addition of 10 μM glibenclamide **(H). (I)** Summary data showing that lowering glucose below 1 mM hyperpolarizes the pericyte membrane, and the effects of 0 glucose were blocked by glibenclamide (4 mM glucose (12 cells, 5 mice) vs. 2 mM glucose (14 cells, 5 mice): *P* > 0.9999, = 0.1204; 4 mM glucose vs. 1 mM glucose (9 cells, 4 mice): *P* = 0.8784, t_97_ = 1.28; 4 mM glucose vs. 750 μM glucose (20 cells, 4 mice): \*\*\**P* = 0.001, t_97_ = 4.04; 4 mM glucose vs. 250 μM glucose (16 cells, 4 mice): \*\*\**P* = 0.0005, t_97_ = 4.193; 4 mM glucose vs. 0 glucose (9 cells, 5 mice): \*\*\**P* = 0.0007, t_97_ = 4.078; 0 glucose vs. 0 glucose + 10 μM glibenclamide (10 cells, 4 mice): \*\*\*\**P* < 0.0001, t_97_ = 5.22; One-way ANOVA with Sidak’s multiple comparison test). **(J)** Concentration-response curve showing pericyte membrane potential hyperpolarizes abruptly in response to lowering glucose concentration.

Under these conditions, activation of K_ATP_ channels with a saturating concentration of pinacidil (10 μM) hyperpolarized Vm by ∼23 mV, an effect that was blocked by the co-application of 10 μM glibenclamide (**Fig. 4B-E**). Strikingly, complete removal of glucose also strongly hyperpolarized the membrane, to -52 mV (**Fig 4G,I**), an effect that was prevented by inclusion of 10 μM glibenclamide in the bath (**Fig. 4 H,I**). Varying glucose within the physiological range measured in the parenchyma (2 mM, 1 mM, 750 μM and 250 μM) revealed the presence of a threshold around 1 mM (EC_50_: 934 μM; **Fig. 4J**), below which a dramatic increase in K_ATP_ channel activity occurs that parallels that seen with 0 glucose, which we refer to as an ‘energy switch’ (**Fig. 4I,J** and **Supplementary Fig. 5**). Together, these data indicate that pericytes monitor small fluctuations of glucose within the physiological range, and if the concentration falls below a critical threshold K_ATP_ channel activity is robustly increased to evoke substantial membrane hyperpolarization.

### GLUT1 block activates the pericyte energy switch *in vivo* and triggers profound arteriolar dilation to increase local blood flow

The endothelium plays a major role in glucose import into the brain, predominately via highly-expressed GLUT1 transporters (**Fig. 5A**), and pericytes also express the gene encoding GLUT1 and to a lesser extent the genes for GLUT3 and GLUT4^20,21^. Given this central role, we hypothesized that blocking GLUT1 would be sufficient to activate the pericyte energy switch and generate K_ATP_ channel activity to hyperpolarize pericyte Vm. This, in turn, should influence electrical signaling through the capillary network and drive an increase in arteriolar diameter and blood flow.

**Figure 5.**
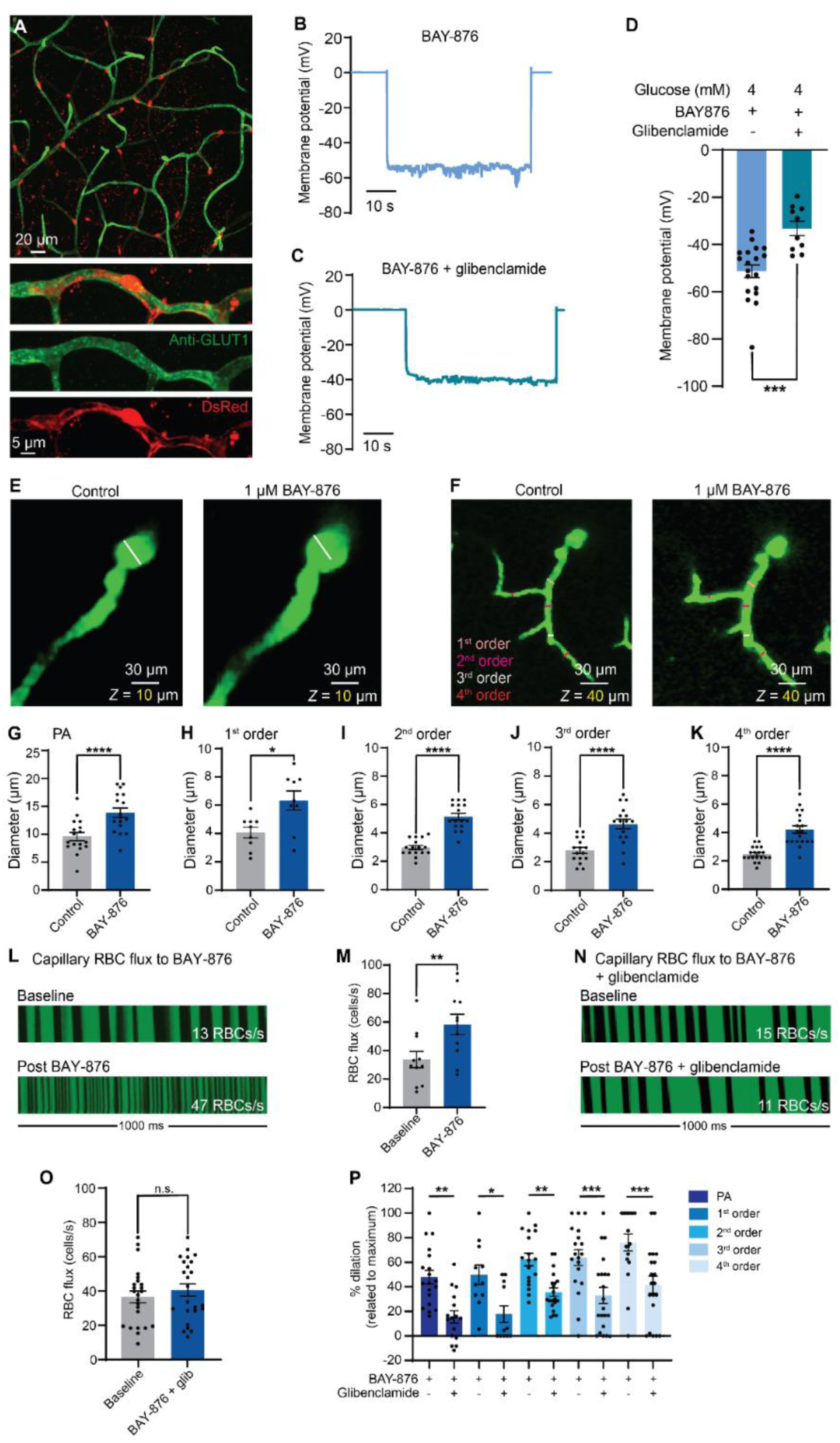
Glucose levels control K_ATP_ channel activity and blood flow *in vivo*. **(A)** Staining with an anti-GLUT1 antibody indicating the high density of this transporter in brain capillaries. **(B-C)** Example traces of membrane potential measurements under 1 μM BAY-876 **(B)** and 1 μM BAY-876 in the presence of 10 μM glibenclamide **(C). (D)** Summary data showing BAY-876 (19 cells, 6 mice) hyperpolarizes pericyte membrane potential and this effect is blocked by glibenclamide (10 cells, 6 mice: \*\*\**P* = 0.0003, t_27_ = 4.192, unpaired Student’s *t*-test). **(E)** Effects of GLUT1 inhibitor BAY-876 (1 μM) on PA diameter. *Left:* PA diameter indicated by white line at baseline. *Right:* Dilation of the same PA after application of BAY-876 to the cranial surface. **(F)** BAY-876 also dilates 1^st^-4^th^ order capillaries. *Left:* A Z-projection showing diameters of 1^st^-4^th^ order at baseline, indicated by colored lines. *Right:* Dilation of the same capillaries after BAY-876 application. **(G-K)** Summary data analyzed using paired Student’s *t*-test, showing dilation across all vessels with BAY-876. **(G)** PA diameter (n = 17 vessels, 5 mice, \*\*\*\**P* < 0.0001, t_16_ = 10.01). **(H)** 1^st^ order capillary diameter (n = 9 vessels, 5 mice, \*\**P* = 0.0029, t_8_ = 4.22). **(I)** 2^nd^ order capillary diameter (n = 16 vessels, 5 mice, \*\*\*\**P* < 0.0001, t_15_ = 11.16). **(J)** 3^rd^ order capillary diameter (n = 16 vessels, 5 mice, \*\*\*\**P* < 0.0001, t_15_ = 8.399) and **(K)** 4^th^ order capillary diameter (n = 17 vessels, 5 mice, \*\*\*\**P* < 0.0001, t_16_ = 7.665). **(L)** Representative 1-s segments of raw kymographs demonstrating hyperemia to BAY-876. *Top*: Baseline RBC flux. *Bottom*: RBC flux measured in the same capillary after BAY-876 application. **(M)** Summary RBC flux data before and after BAY-876 application (n = 11 paired measurements, 4 mice, \*\**P* = 0.004, t_10_ = 3.71, paired Student’s *t*-test). **(N)** The blood flow response to BAY-876 is mediated by K_ATP_ channel activation. *Top:* RBC flux at baseline. *Bottom:* RBC flux measured from the same capillary showing no change in blood flow after the application of BAY-876 in the presence of K_ATP_ channel blocker glibenclamide (10 μM). **(O)** Summary RBC flux data from >5^th^ order capillaries when BAY-876 was applied in the presence of glibenclamide (n = 24 paired measurements, 5 mice, *P* = 0.2324, t_23_ = 1.226, paired Student’s *t*-test). **(P)** Summary data showing significantly decreased dilatory responses to BAY-876 in the presence of glibenclamide across all vessel orders (n = 5 mice per group; PA: \*\**P* = 0.0013, t_163_ = 3.734; 1^st^ order capillary: \**P* = 0.0189, t_163_ = 2.936; 2^nd^ order capillary: \*\**P* = 0.0079, t_163_ = 3.212; 3^rd^ order capillary: \*\*\**P* = 0.0009, t_163_ = 3.825; 4^th^ order capillary: \*\*\**P* = 0.0003, t_163_ = 4.14; one-way ANOVA with Sidak’s multiple comparison test).

In line with the predictions of our hypothesis, blocking glucose entry using the selective GLUT1 inhibitor BAY-876 (1 μM) hyperpolarized the pericyte membrane to -51 mV, as seen with concentrations of glucose below 1 mM (**Fig. 4I**), and this effect was completely inhibited by glibenclamide (**Fig. 5B-D**). Based on the known Vm-diameter relationship of PA smooth muscle, a ∼15-mV hyperpolarization is predicted to dilate PAs by approximately 50% (see ref 43). Accordingly, we tested the effect of 1 μM BAY-876 on PA and capillary diameter, and capillary blood flow when applied directly to the cranial surface *in vivo*. Strikingly, this maneuver produced a 48% increase in PA diameter (**Fig. 5E,G**), in line with our predictions, and profoundly dilated 1^st^-4^th^ order capillaries (**Fig. 5F,H-K**) while also almost doubling capillary blood flow (**Fig. 5L,M**). Pre-incubation with glibenclamide (10 μM) eliminated the BAY-876–evoked increase in capillary RBC flux (**Fig. 5N,O**) and significantly decreased the dilatory effect of BAY-876 at the level of the PA (68% reduction) and in 1^st^-4^th^ order capillaries (**Fig. 5P**). Together, these data indicate that a reduction in glucose delivery to the pericyte interior triggers K_ATP_ channel-mediated electrical signaling, which in turn is transmitted upstream to the PA to drive dilation and an increase blood flow.

## DISCUSSION

Taken together, our data reveal that K_ATP_ channels in capillary thin-strand pericytes couple changes in energy substrate levels to alterations of local brain blood flow. Our data support a model in which pericyte K_ATP_ channels initiate robust hyperpolarization in response to a decrease in local glucose below a critical threshold, which can be transferred over long distances through engagement of capillary electrical signaling, eliciting relaxation of remote arteriolar SMCs, leading to vasodilation and an increase in blood flow into the capillary bed (**Fig. 6**).

**Figure 6.**
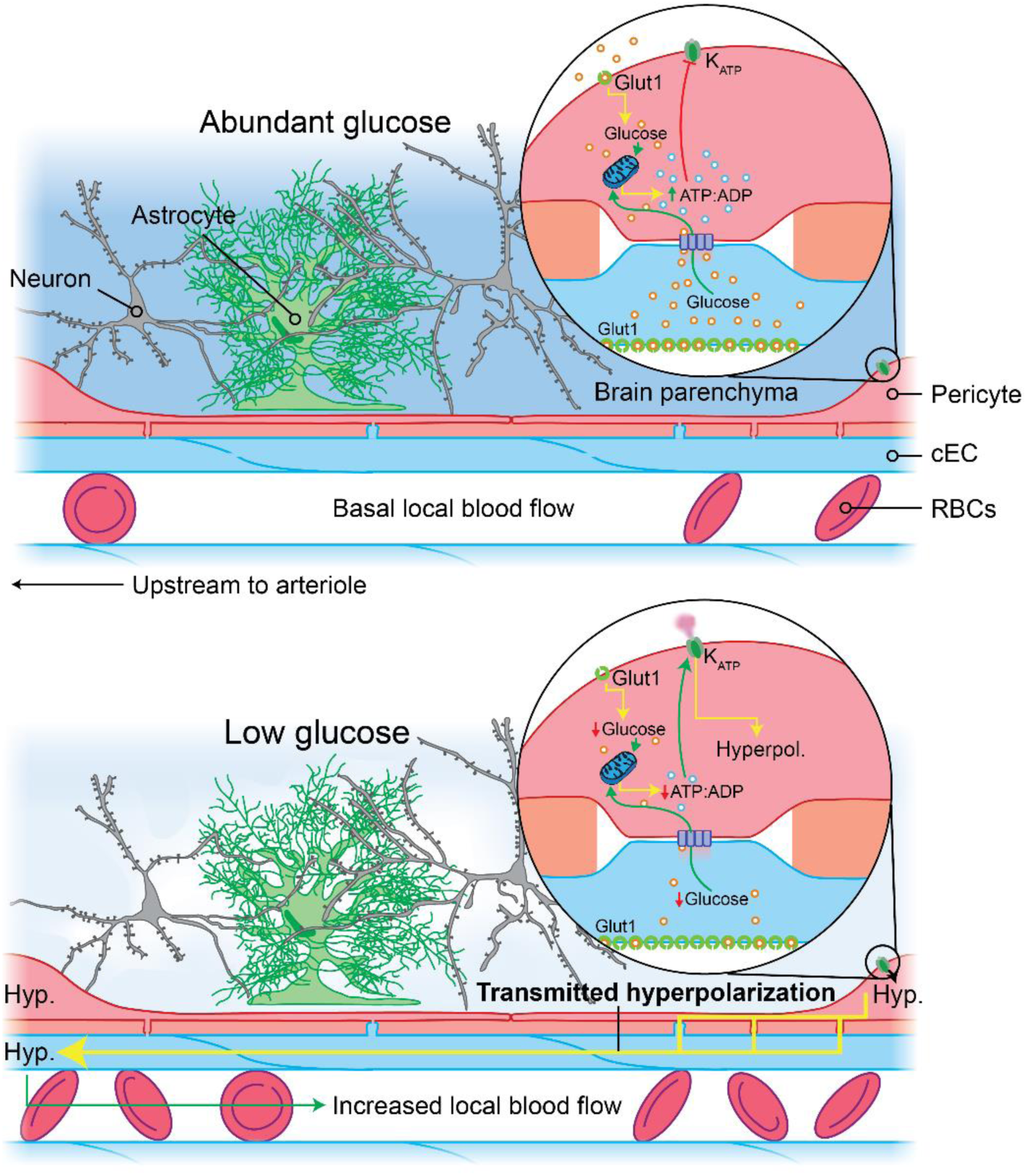
Illustrative summary and model for pericyte K_ATP_ channel-mediated coupling of electrical activity with glucose availability. Under conditions of abundant glucose, pericyte ATP:ADP ratio is high and keeps pericyte K_ATP_ channels closed. A decrease in GLUT-1 mediated glucose import, or a drop in local glucose availability results in pericyte K_ATP_ channel activation, likely due to a corresponding decrease in cellular ATP:ADP ratio. When activated, K_ATP_ channels robustly hyperpolarize pericyte membrane potential and this electrical signal is then fed into the underlying capillary endothelium to be rapidly transmitted upstream via a K_IR_2.1 channel-dependent mechanism. This remotely dilates penetrating arterioles and increases blood flow, thereby replenishing local glucose levels and protecting ongoing neuronal metabolism and function.

### A pericyte energy switch: membrane potential is steeply influenced by local glucose concentration

The brain relies primarily on glucose and oxygen to fuel its energy requirements. The central pathway for glucose entry into the brain is via the GLUT1 transporter, which is abundantly expressed in blood brain barrier ECs^21,44^. The cell bodies and processes of pericytes that decorate the vascular wall are embedded in the basement membrane that surrounds capillary ECs, and this intimate association allows for the extension and receipt of projections known as peg-socket junctions^11^ which bring the membranes of these two cell types into very close proximity and likely facilitates the formation of gap junctions^10,19,27^. Given that gap junctions permit transfer of molecules up to 1000 Da, combined with observations of cell-cell transfer of fluorescently-conjugated glucose analogues^45,46^, it is reasonable to posit that glucose (∼180 Da) taken up into the EC cytoplasm may be transferred directly to pericytes via this avenue, the rate of which will depend ultimately on the degree of coupling between these cell types. Pericytes also express several GLUT-encoding genes (*Slc2a1* > *Slc2a4* > *Slc2a3*, which translate to GLUTs 1, 4, and 3, respectively^20,21^), suggesting that they may also be capable of taking up glucose directly from their surroundings. Collectively, these molecular features likely equip capillary pericytes to sense and closely monitor glucose levels in their locale.

As a result, we hypothesized that pericytes may be capable of responding to changes in local glucose availability through metabolically-evoked K^+^ channel activity and blood flow modulation, by virtue of their robust K_ATP_ channel expression. Thus, to directly ascertain whether pericyte electrical behavior is influenced by local energy availability, we sought to determine the relationship between glucose concentration and pericyte Vm in granular detail, and specifically focused on the contribution of K_ATP_ channels in this context. Accordingly, we tested the effects of lowering glucose from 4 mM (the concentration typically found in bulk CSF) to 1 mM and below (which aligns with measurements several independent groups have made of parenchymal glucose concentrations^34–42^). We found that complete removal of glucose produced a striking 16-mV hyperpolarization, mediated by K_ATP_ channel activation. Moreover, almost identical responses were seen for glucose concentrations up to 0.75 mM and in circumstances in which we blocked glucose import via GLUT1. In contrast, 2 mM glucose had little influence on Vm, which remained close to the ‘resting’ value we obtained in 4 mM glucose (−36 mV). At 1 mM glucose, pericyte Vm was slightly more hyperpolarized (−40 mV) but this was not significantly different than higher glucose concentrations. These data indicate that pericytes are steeply sensitive to local changes in this key energy substrate, and are consistent with the existence of a glucose concentration threshold below which robust activation of K^+^ efflux through K_ATP_ is elicited. This ‘all-or-none’ effect of energy substrate abundance on membrane potential—reminiscent of flipping a switch—may be triggered by changes in glucose affecting the production of ATP in the pericyte, leading to a new set point for the intracellular ATP:ADP ratio. Accompanying this could be an amplification mechanism such as the engagement of capillary K_IR_ channels, which are directly activated by membrane hyperpolarization relieving voltage-dependent block of the channel pore by polyamines^47^. K_IR_ channel activation in turn may boost K_ATP_-initiated hyperpolarization and combined, these factors could translate a change in intracellular metabolism into a binary response, driving Vm towards E_K_ and facilitating potent hyperemic responses to small changes in external glucose availability. Alternatively, or perhaps in conjunction, other energy-sensing molecules such as adenosine monophosphate-activated protein kinase (AMPK) may be engaged by glucose deficits to phosphorylate K_V_ channels (which have been reported in cultured retinal pericytes but await confirmation in native cells^48^) and increase their activity^49^. As cECs have also recently been shown to possess K_ATP_ channels, albeit at lower current density^25^, we cannot presently fully rule out the possibility of their contribution to these K_ATP_ channel-mediated effects on Vm, although our imaging data are consistent with pericytes playing the major role. Further experiments are needed to explore these possibilities in detail.

What might be the circumstances, physiological and pathological, that engage this mechanism? One possibility is that local fluctuations in glucose that occur during concerted neuronal activity^50,51^ continually adjust the electrical input of pericytes to the capillary endothelium, resulting in fine-tuning of local blood flow to ensure that neuronal metabolism is protected on a moment-to-moment time scale. Given that K_ATP_ channels do not appear to contribute to functional hyperemia to a diffuse visual stimulus^52^, it may be that strong stimuli driving robust network activity and rapidly ramping energy demands are required to engage this mechanism under physiological conditions. It is also possible that the pericyte energy switch is reserved for pathological conditions such as hypoglycemia, a common occurrence in diabetic individuals, where it might serve as an emergency failsafe that has evolved to protect brain energy supply by increasing blood flow. In support of these ideas, as parenchymal glucose approaches 0, blood flow has been observed to increase by up to 57% (ref 35), and insulin-induced hypoglycemia increases blood flow by 42% in adults^53^.

### Pericyte K_ATP_ channel-meditated electrical signals are transmitted through multiple branch orders to control blood flow

Thin-strand pericytes are found deep within the capillary bed, starting around the 5^th^ order branches and above. An elegant recent study deploying optogenetic tools in pericytes has shown that these cells are capable of exerting slow constrictions of their underlying capillaries^6^, yet it appears that they do not dilate during functional hyperemia^16,54^. To rapidly control blood flow, thin-strand pericytes could modulate ongoing electrical signaling through the underlying endothelium which is transmitted over long distances to influence upstream arteriolar diameter^28^. Together, the present experiments support this idea and reveal that focal activation of the K_ATP_ channels in just a single pericyte is sufficient to evoke rapid dilation of remote PAs at distances up to at least 421 μm (the furthest site we stimulated in our experiments). In stark contrast, we did not detect an effect of direct stimulation of PAs with 10 μM pinacidil on diameter, although we note that application of a 500-fold higher concentration onto PAs and 1^st^-3^rd^ order capillaries caused localized vasodilation in another study^17^. Our data using lower (but saturating) concentrations of pinacidil suggest that functional K_ATP_ channels are absent, or present at too low of a density in the arteriolar wall to generate sufficient hyperpolarization to elicit vasorelaxation under the conditions used here, and this is consistent with previous findings in isolated and pressurized PAs which did not dilate to bath application of a K_ATP_ channel activator^26^. Moreover, pinacidil-stimulation of ECs on a segment of 5^th^ or higher order capillary lacking a pericyte cell body, or of pericytes with genetically inactivated K_ATP_ channels, failed to dilate PAs. Collectively, these observations strongly imply that pericytes represent the locus of K_ATP_ channel activity in the capillary bed, and from this locus, hyperpolarization must then be transmitted to upstream SMCs to evoke dilation at a distance. There are several possibilities as to how such long-range communication may be achieved. The eponymous projections of thin-strand pericytes reach over long distances and come into close contact with those of neighboring cells. However, these do not appear to closely interdigitate and rather stay confined to their own territories^8^, and no evidence of direct pericyte-pericyte transfer of charge or chemical agents has been reported to our knowledge, with the exception of specialized interpericyte tunneling nanotube (IPNT) projections in the retina^55^. Thus, it presently seems unlikely that capillary pericytes without IPNTs directly exchange electrical signals. Instead, mounting evidence indicates that thin-strand pericytes directly interface with capillary ECs via gap junctions^10^. Our prior work^28^ revealed that electrical signaling through the brains capillary network to upstream arterioles is a major mechanism for blood flow control in the brain. This mechanism relies on capillary EC K_IR_2.1 channels, which are activated by both external K^+^ and membrane hyperpolarization and transmit electrical signals upstream at a velocity of several millimeters per second^23,28^. Given that our data show that the K_IR_2 channel blocker Ba^2+^ eliminates pinacidil-evoked remote dilation of PAs, our observations in context with those of other groups cumulatively suggest that the activation of K_ATP_ channels in pericytes generates membrane hyperpolarization that is then injected via peg-socket junctions into the underlying ECs to engage capillary EC electrical signaling and dilate upstream arterioles. We also recently reported that capillary EC Ca^2+^ signals control blood flow through a nitric oxide-dependent mechanism that relaxes contractile pericytes of the 1^st^ to 4^th^ order transitional segment of the capillary bed^18^. Intriguingly, these signals are strongly influenced by ongoing electrical signaling in the capillaries, with the hyperpolarization these provide likely increasing the driving force for Ca^2+^ entry. Thus, it is possible that pericyte K_ATP_-mediated electrical signals might also promote capillary EC Ca^2+^ signaling, which could also be a contributory factor in the observed dilations resulting from these.

### Recasting Pericytes as Metabolic Sentinels

Pericytes play a range of roles in the brain, which include control of blood-brain barrier function^12^, regulation of endothelial gene expression^13^, promotion of proper vascular development^56^, provision of structural stability^57^, and regulation of blood flow^15,58^. Moreover, they appear to be particularly vulnerable cells in the context of dementias and a range of other disorders impinging on brain function (e.g. diabetes, hypertension and kidney dysfunction^59^), and contractile pericytes have been noted to die in rigor which is thought to contribute to loss of brain blood flow control, ultimately precipitating neuronal dysfunction and decline^15^. Our data evoke novel concepts stemming from the sensitivity of thin-strand pericytes to subtle metabolic changes. Importantly, if glucose drops below a critical threshold, a robust electrical response is generated through the recruitment of pericyte K_ATP_ channels to increase local blood flow, thereby providing more glucose to replenish local levels and protect ongoing neuronal function. This mechanism may be critical for the maintenance of brain health, and its disruption over long periods could contribute to the mismatch between energy supply and demand that occurs in cognitive decline and dementia^60– 62^. Intriguingly, a recent VINE-seq atlas of human vascular cells suggest that the molecular players that take center-stage in the electrical switch we have elucidated here (*Kcnj8, Kcnj2*, and *Slc2a1*) are each profoundly downregulated in Alzheimer’s cerebrovasculature, which could potentially disable protective responses to local glucose dips and imperil neuronal metabolism^63^. In support of this idea, Kir2.1 function is known to be disrupted in the 5xFAD mouse model of AD^64^. Further work is needed to address whether the pericyte energy switch is disabled in Alzheimer’s, and ongoing experiments in our laboratory are now directly addressing these questions.

Our observation that pericytes are sensitive to glucose naturally evokes the question of whether pericytes detect the levels of other energy substrates and metabolites. Pericytes exist in, and are influenced by, a rich milieu of molecules and substrates, of which glucose is just one element. Therefore, pericyte activity is likely to be regulated by a complex mix of factors which fluctuate in concentration over widely varying timescales. One such factor, partnered with glucose to support brain metabolism, might be oxygen. Oxidative phosphorylation relies on local oxygen tension, which in turn is a direct function of local blood flow^65^. The oxidation of glucose provides vastly more ATP than glycolysis alone and neuronal activity is primarily powered by oxidative phosphorylation^66^. It is possible that the pericyte energy switch may also be activated by local transient decreases in oxygen^67^, which might lead to an abrupt fall in intracellular ATP production, influencing ATP:ADP ratio and engaging K_ATP_ channels. Interestingly, stalling (i.e. complete cessation of RBC flux) behavior is relatively common in brain capillaries, with ∼0.45% of capillaries estimated to be stalled at any one time^68^. The function of this phenomenon is unclear, but it seems likely that these events would lead to a localized decrease in oxygen tension due to the lack of transiting RBCs loaded with oxygen. This may in turn activate the pericyte energy switch, leading to signaling to increase blood flow to relieve the stall before it damages neurons. Still other metabolites might be sensed by pericytes and evoke K_ATP_-mediated hyperpolarizing responses. As we previously noted^19^ pericytes express the A_2a_ adenosine receptor, a G_s_-coupled GPCR, activation of which has recently been shown to lead to pericyte K_ATP_ channel activation through protein kinase A^25^. As adenosine is released from neurons during their activity, this pathway may also engage pericyte K_ATP_ channel activity to hyperpolarize the cell membrane and evoke upstream arteriolar dilation as we have shown here. Pericytes might also possess mechanisms to assess local carbon dioxide gradients^69^, which would reflect the degree of local metabolic activity^70^, and may modulate blood flow in turn. It has also recently been demonstrated that pericytes can sense lactate generated during glycolysis in ECs^71^, which could also serve as an energy substrate that ultimately regulates pericyte K_ATP_ channel activity.

## SUMMARY AND CONCLUSION

Despite their intimate association with capillaries, the precise contribution of thin-strand pericytes to the control of blood flow in the brain is largely unknown. A rich complement of ion channels and GPCRs equips pericytes to sense and respond to a wide range of stimuli^19^. K_ATP_ channels are the most abundant ion channel expressed by pericytes^19^, and we demonstrate here that their activation in response to decreased local metabolic substrate availability produces a robust increase in blood flow. Our data thus recast pericytes as metabolic sentinels that form a brain-wide energy-sensing network, continually monitoring glucose concentrations and adjusting blood flow to protect ongoing neuronal health and function. In conditions like sporadic Alzheimer’s disease, brain glucose levels and metabolism are profoundly dysregulated^72–75^ and thus determining the impact of this on pericyte energy sensing and accompanying blood flow control may yield potential targets for improving clinical outcomes in neurological diseases with a significant vascular and metabolic component.

## METHODS

### Animal husbandry

Adult (2–3 mo. old) male and female C57BL/6J mice, *Cspg4*-DsRed mice (C57BL/6J background; Jackson Laboratories), *Cspg4*-Cre recombinase mice, and *Cspg4*-Cre-K_IR_6.1^AAA^ mice were group-housed on a 12-h light:dark cycle with environmental enrichment and free access to food and water. Tamoxifen inducible *Cspg4*-Cre-K_IR_6.1^AAA^ mice were generated by crossing K_IR_6.1^AAA^ mice expressing dominant-negative K_IR_6.1^AAA^ with *Cspg4*-Cre recombinase mice^29,30^. All animal procedures received prior approval from the University of Maryland Institutional Animal Care and Use Committee.

### K_IR_6.1^AAA^ induction

4-OHT, the active metabolite of tamoxifen, was dissolved in a corn oil:ethanol solution (90:10% v/v) at a concentration of 2 mg/ml ^76^. *Cspg4*-Cre-K_IR_6.1^AAA^ mice were given either 4-OHT (10 mg/kg, intraperitoneal; K_IR_6.1^AAA^ induction) or vehicle (corn oil:ethanol; vehicle control), and control K_IR_6.1^AAA^ mice were given 4-OHT (10mg/kg, intraperitoneal; Cre control) once a day for 5 consecutive days. 4 weeks after the last injection, mice were imaged *in vivo* as described below.

### Chemicals

BAY-876 was purchased from Tocris Bioscience (USA). All other chemicals were obtained from Sigma Aldrich (USA).

### *In vivo* imaging

Cranial window preparation and *in vivo* imaging was performed as previously described^18,28^. Briefly, mice were anesthetized with isoflurane (5% induction, 1.5-2% maintenance). 150 μL of FITC-dextran (10mg/ml) or TRITC-dextran (40 mg/ml) dissolved in saline was injected retro-orbitally. A midline incision was made on the scalp to expose the skull, and a titanium head plate was affixed over the left hemisphere with a combination of dental cement and superglue. On securing the headplate in a holding frame, a circular cranial window (∼2 mm diameter) was drilled in the skull over the somatosensory cortex. The skull piece was removed, and the dura was carefully resected. The cranial surface was irrigated as necessary with saline. Upon conclusion of surgery, isoflurane anesthesia was replaced with α-chloralose (50 mg/kg) and urethane (750 mg/kg). Body temperature was maintained at 37°C throughout the experiment using a rectal probe feedback-controlled electric heating pad (Harvard Apparatus). Oxygenated and warmed (35-36 °C) aCSF (124 mM NaCl, 3 mM KCl, 2 mM CaCl_2_, 2 mM MgCl_2_, 1.25 mM NaH_2_PO_4_, 26 mM NaHCO_3_, 4 mM glucose) was superfused over the exposed cortex for the duration of the experiment at a rate of ∼1 mL/min and continuously monitored at the window with a temperature probe. Images were acquired through an Olympus 20x infinity-corrected Plan Fluorite 1.0 NA water-immersion objective mounted on a Scientifica Hyperscope (Scientifica, UK) coupled to a Coherent Chameleon Vision II Titanium-Sapphire pulsed fs laser (Coherent, USA). FITC- and TRITC-dextran or DsRed were excited at 820 nm or 920 nm, respectively, and emitted fluorescence was separated through 525/50 and 620/60 nm bandpass filters. Single-plane imaging data to examine the time course of vessel diameter changes was collected at 30 Hz using a resonant scanning mirror. 3D imaging data were typically gathered using standard galvo mirrors. To measure RBC flux, we performed line scans at 1 kHz. Line scans were oriented along the lumen parallel to the flow of blood to maximize flux signal. For pressure-ejection of agents in aCSF (vehicle) onto pericytes or endothelial cells, a pipette containing the agent of interest and FITC or TRITC-dextran (to enable visualization) was maneuvered into the cortex and positioned adjacent to the cell under study, after which the solution was ejected directly at 8–12 psi, for 30 ms. This approach restricted agent delivery to the target cell and caused minimal displacement of the surrounding tissue. For pharmacological and staining experiments, agents of interest were applied to the cranial surface for a minimum of 20 min to allow penetration. All *in vivo* imaging experiments were routinely ended with the application of aCSF containing 0 Ca^2+^ supplemented with 5 mM EGTA and 200 μM diltiazem to elicit maximal relaxation of SMCs and contractile pericytes to enable the measurement of maximum vessel diameters.

### Microelectrode impalement of pericytes on isolated microvessels

Membrane potential measurements were made by impaling pericytes on microvessels isolated from *Cspg4*-DsRed mice using a papain-based Neural Tissue Dissociation kit (Miltenyi Biotec), as described previously^21,25^. Cortical tissue from one hemisphere was carefully dissected and minced into small pieces with microscalpels in an isolation solution containing 55 mM NaCl, 80 mM Na-glutamate, 5.6 mM KCl, 2 mM MgCl_2_, 10 mM HEPES and 4 mM glucose (pH 7.3). Minced tissue was incubated with enzyme P from the kit for 18 min at 37°C, followed by addition of enzyme A, homogenization by passing through a Pasteur pipette ∼10 times and incubation for 15 min at 37°C. The homogenate was then passed through a 21 G needle 7 times and incubated for 12 min at 37°C. The cell suspension was filtered through a 62-μm nylon mesh and stored in ice-cold isolation solution. Cells were transferred to a silicone elastomer (SYLGARD 182)-lined perfusion chamber, and allowed to adhere for ∼45 min. The chamber was perfused with bath solution consisting of 137 mM NaCl, 3 mM KCl, 2 mM CaCl_2_, 1 mM MgCl_2_, 10 mM HEPES and 0-4 mM glucose (pH 7.4). DsRed-positive pericytes on small capillary segments were identified using brightfield microscopy, and impaled with a sharp microelectrode (pulled to ∼100-200 MΩ) filled with 0.5 M KCl. Only recordings fulfilling the following criteria were considered for analysis: stable baseline prior to impalement, sharp negative deflection of membrane potential upon impalement, immediate return to 0 mV upon withdrawing the electrode. Membrane potential was recorded using an AxoClamp 900A digital amplifier and HS-2 headstage (Molecular Devices). Signals were digitized and stored using Axon Digidata 1550B and pClamp 9 software (Molecular Devices).

### Immunohistochemistry

Brains were extracted from *Cspg4*-DsRed mice that underwent cardiac perfusion with 4% paraformaldehyde. Tissues were stored in 4% paraformaldehyde overnight at 2–8ºC and dehydrated in 30% sucrose in 1x phosphate buffered saline (PBS). Immunostaining and optical clearing of brain samples were performed according to a modified CUBIC clearing method^77,78^. Briefly, fixed brains were immunostained by first blocking non-specific binding with normal goat serum (Vector Laboratories, USA). Blocked samples were incubated overnight at 2–8ºC with rabbit anti-SLC2A1 polyclonal antibody (1:500 dilution, HPA031345, Atlas Antibodies, Stockholm, Sweden) and developed with Alexa Fluor 488 goat anti-human IgG secondary antibody (1:1000). Samples were cleared by incubation in CUBIC R1 solution (see ref 77) at 37°C with shaking for 2-3 weeks, and then incubated in RIMS (refractive index matching solution; 88% w/v Histodenz in 0.02 M PBS with 0.01% sodium azide) at 37°C until the samples were optically clear (∼5 days) with solution being replaced every 24 hours. Cleared tissue was mounted in RIMS and imaged with a Nikon W1 spinning disk confocal microscope.

### Data analysis and statistical testing

Diameter measurements were analyzed offline using ImageJ software. Vessel diameter was calculated as the average of three measurements per vessel type made from Z-stacks of 3D volume recordings using the full-width at half-maximum method. RBC flux data were binned at 1-s intervals and analyzed using SparkAn software (A. Bonev, University of Vermont). For pressure-ejection experiments, mean baseline diameter and flux were obtained by averaging the baseline for each measurement before ejection of pinacidil or aCSF, and peak diameter and RBC flux change was defined as the largest change from mean baseline. The distance from the site of pressure ejection to the feed arteriole was estimated using the Simple Neurite Tracer plugin on ImageJ software^79^. Statistical testing was performed using GraphPad Prism 7 software. Data are expressed as means ± s.e.m., and a *P*-value ≤ 0.05 was considered significant. Stars denote significant differences; ‘n.s.’ indicates comparisons that did not achieve statistical significance. Statistical tests are noted in figure legends. All t-tests were two-sided. Statistical methods were not used to pre-determine sample sizes, and ample size was estimated based on similar experiments performed previously in our laboratory. Experiments were repeated to adequately reduce confidence intervals and avoid errors in statistical testing. Data collection was not performed blinded to the conditions of the experiments. Littermates were randomly assigned to experimental groups; no further randomization was performed. No data were excluded.

## ACKNOWLEDGEMENTS

The authors thank B. Huang and S. Edwards for animal husbandry and experimental support. Support for this work was provided by the NIH National Institute on Aging and National Institute of Neurological Disorders and Stroke (1R01AG066645, 5R01NS115401, and 1DP2NS121347-01, to T.A.L), and the American Heart Association and the D.C. Women’s Board (Award 830093 to A.H,17SDG33670237 and 19IPLOI34660108 to T.A.L).

## AUTHOR CONTRIBUTIONS

A.H. designed experiments, acquired and analyzed data, and edited the manuscript. C.R acquired and analyzed pinacidil surface application data, D.G performed immunofluorescence and imaging of GLUT1 staining, T.A.L directed the study, acquired and analyzed data, and edited the manuscript. All authors reviewed the manuscript and approved its submission.

## SUPPLEMENTAL INFORMATION

**Supplementary figure 1.**
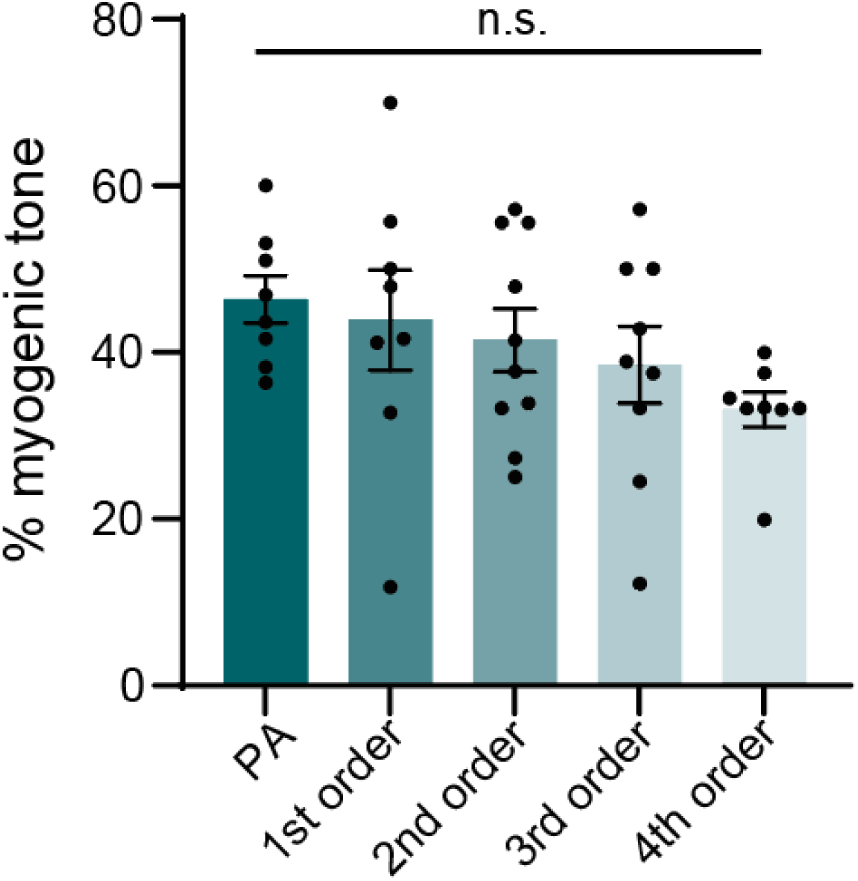
Summary data of myogenic tone of PAs and capillaries *in vivo*. PA (8 vessels, 5 mice) vs. 1^st^ order capillary (8 vessels, 5 mice): *P* = 0.9814, q_38_ = 0.41; PA vs. 2^nd^ order capillary (10 vessels, 5 mice): *P* = 0.8037, q_38_ = 0.8518; PA vs. 3^rd^ order capillary (9 vessels, 5 mice): *P* = 0.4761, q_38_ = 1.336; PA vs. 4^th^ order capillary (8 vessels, 5 mice): *P* = 0.1092, q_38_ = 2.185; one-way ANOVA with Dunnett’s multiple comparison test.

**Supplementary figure 2.**
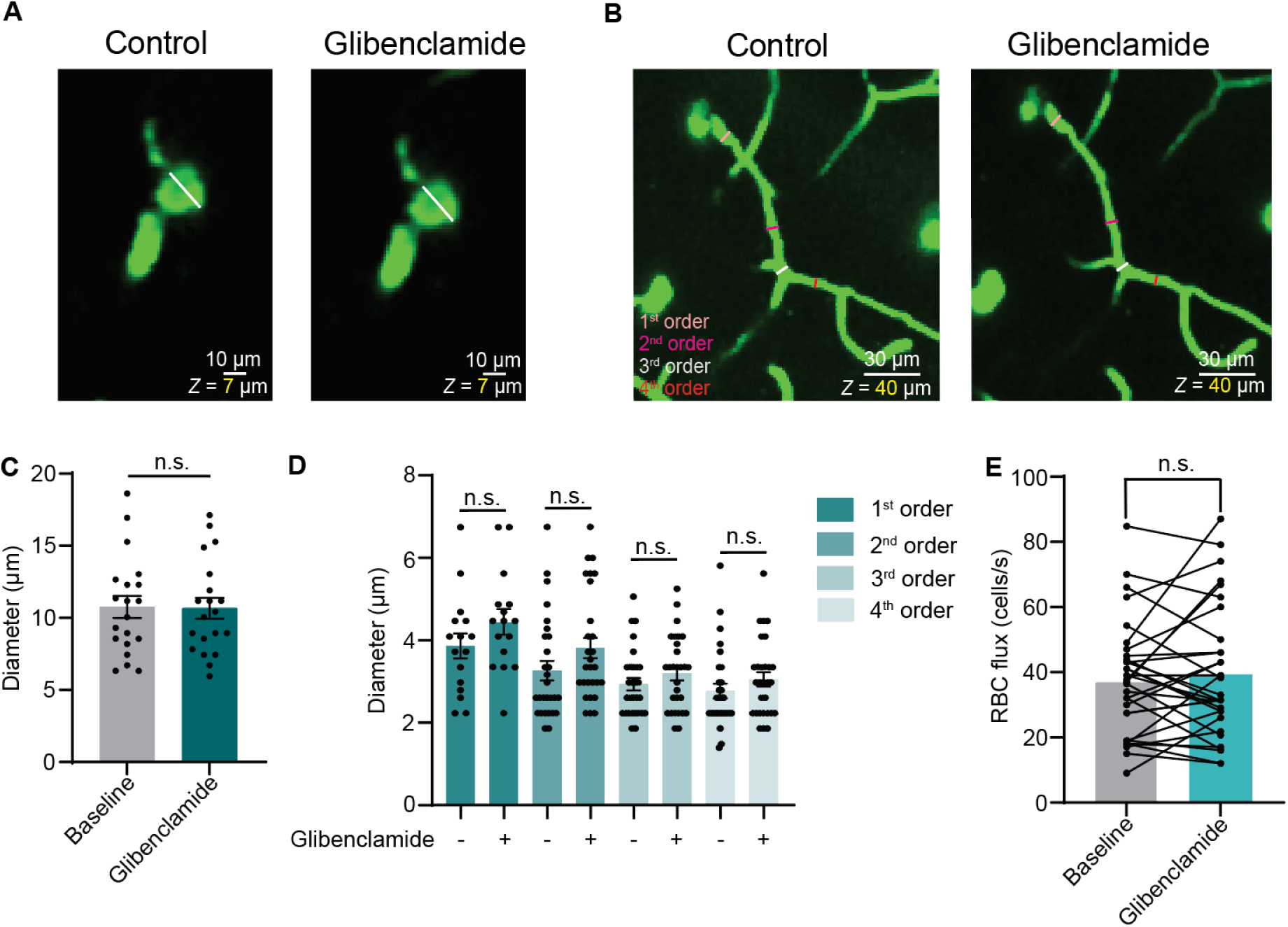
Vascular K_ATP_ channel activity is minimal at rest *in vivo*. **(A)** Effects of glibenclamide (10 μM) on PA diameter. *Left*: PA diameter indicated by white line at baseline. *Right:* The same PA after application of glibenclamide. **(B)** A PA and its downstream 1^st^-4^th^ order capillaries. *Left:* Baseline diameters of 1^st^-4^th^ order capillaries indicated by respective colored lines. *Right:* The same 1^st^-4^th^ order capillaries after application of glibenclamide. **(C)** Summary data of PA diameter before and after application of glibenclamide (n = 20 paired measurements, 6 mice, *P* = 0.7779, t_19_ = 0.2861, paired Student’s *t*-test). **(D)** Summary data of 1^st^ – 4^th^ order capillary diameter showing no change after glibenclamide application (n = 6 mice per group; 1^st^ order capillary: *P* = 0.4326, t_196_ = 1.512; 2^nd^ order capillary: *P* = 0.2001, t_196_ = 1.936; 3^rd^ order capillary: *P* = 0.8198, t_196_ = 0.9398; 4^th^ order capillary: *P* = 0.7720, t_196_ = 1.020; one-way ANOVA with Sidak’s multiple comparison test). (E) Summary RBC flux data from >5^th^ order capillaries demonstrating no change in blood flow after application of glibenclamide (n = 32 paired measurements, 6 mice, *P* = 0.4487, t_31_ = 0.7674, paired Student’s *t*-test).

**Supplementary figure 3.**
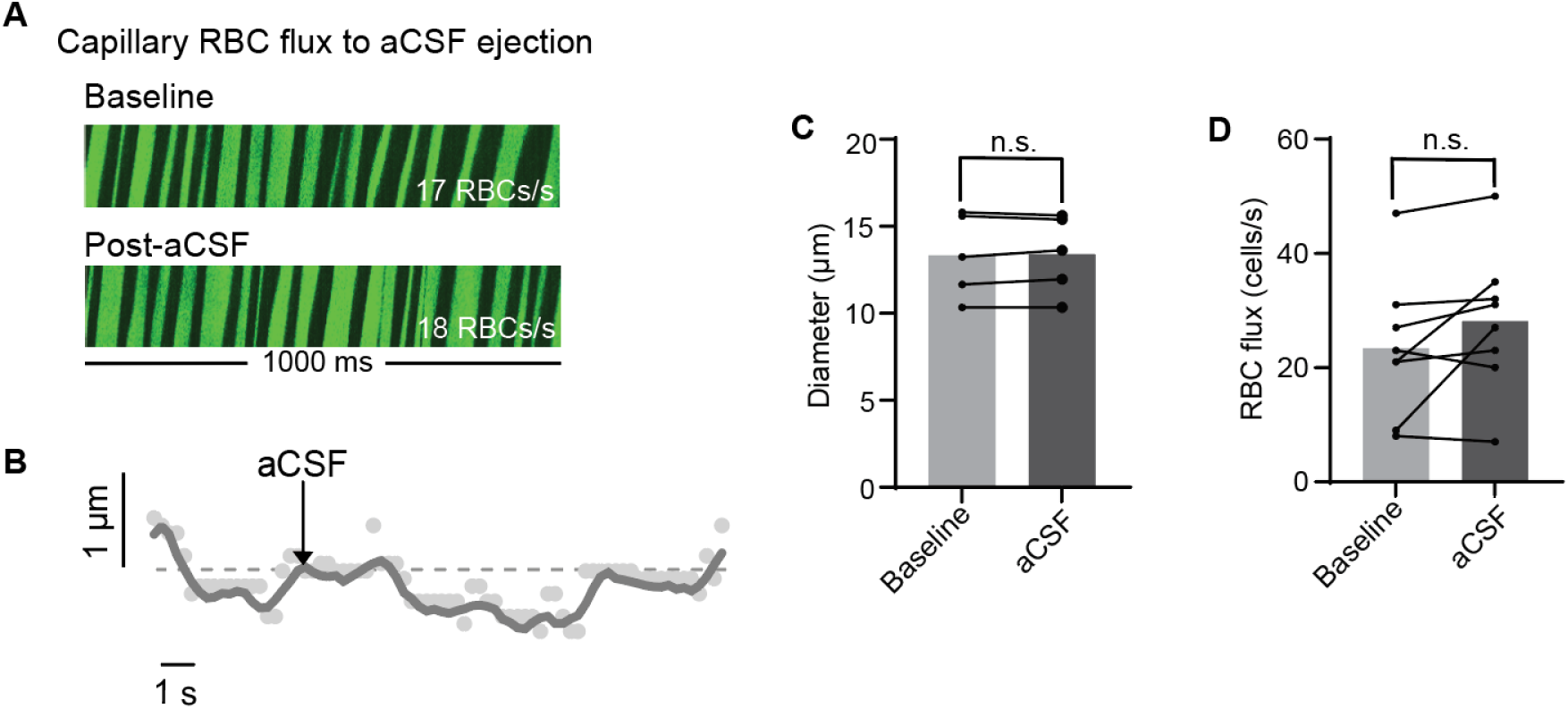
Direct stimulation of a pericyte with vehicle (aCSF) does not dilate PAs or increase blood flow. **(A)** 1-s kymograph segments showing raw RBC flux of a >5^th^ order capillary at baseline, and after aCSF was ejected onto the overlying pericyte. **(B)** Representative time course showing no change PA diameter after direct stimulation of a pericyte with aCSF. **(C)** Summary data showing PA diameter before and after ejection of aCSF on a pericyte (n = 5 paired measurements, 4 mice, *P* = 0.6317, t_4_ = 0.5181, paired Student’s *t*-test). **(D)** Summary data of RBC flux before and after aCSF-ejection on a pericyte (n = 8 paired measurements, 4 mice, *P* = 0.1108, t_7_ = 1.825, paired Student’s *t*-test).

**Supplementary figure 4.**
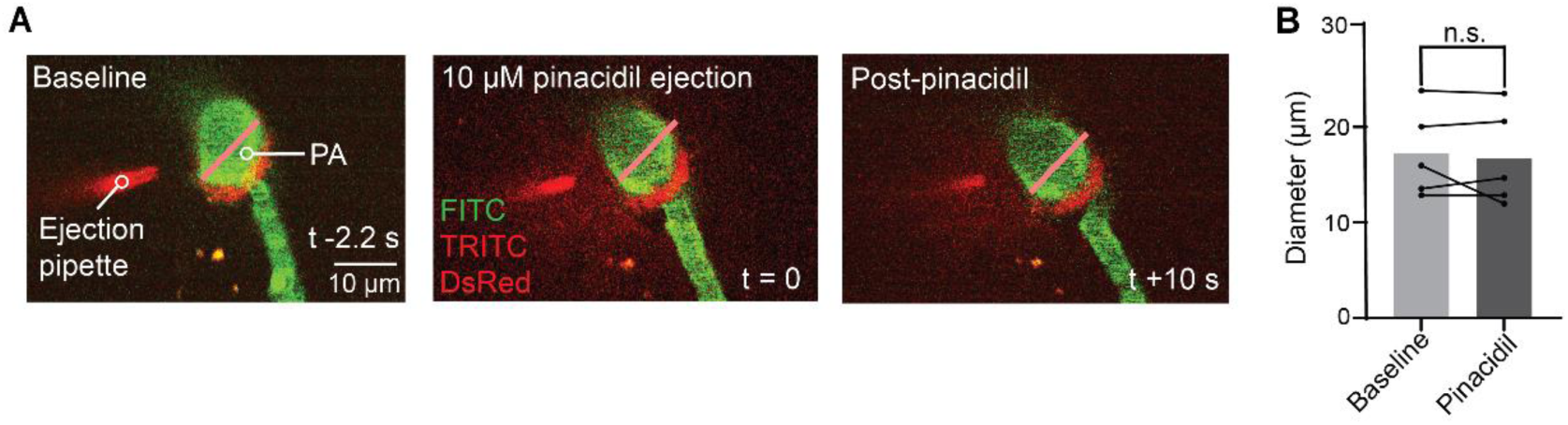
Ejection of pinacidil on a PA does not affect its diameter **(A)** Focal stimulation of a PA with 10 μM pinacidil. *Left*: PA diameter at baseline indicated by pink line, and an ejection pipette containing 10 μM pinacidil positioned next to the PA. *Middle*: Ejection of pinacidil directly onto the PA. *Right*: PA diameter 10 s after pinacidil ejection compared to control diameter, indicated by pink line. **(B)** Summary data showing no change in PA diameter after direct stimulation with pinacidil (n = 5 paired measurements, 4 mice, *P* = 0.5946, t_4_ = 0.5774, paired Student’s *t*-test).

**Supplementary figure 5.**
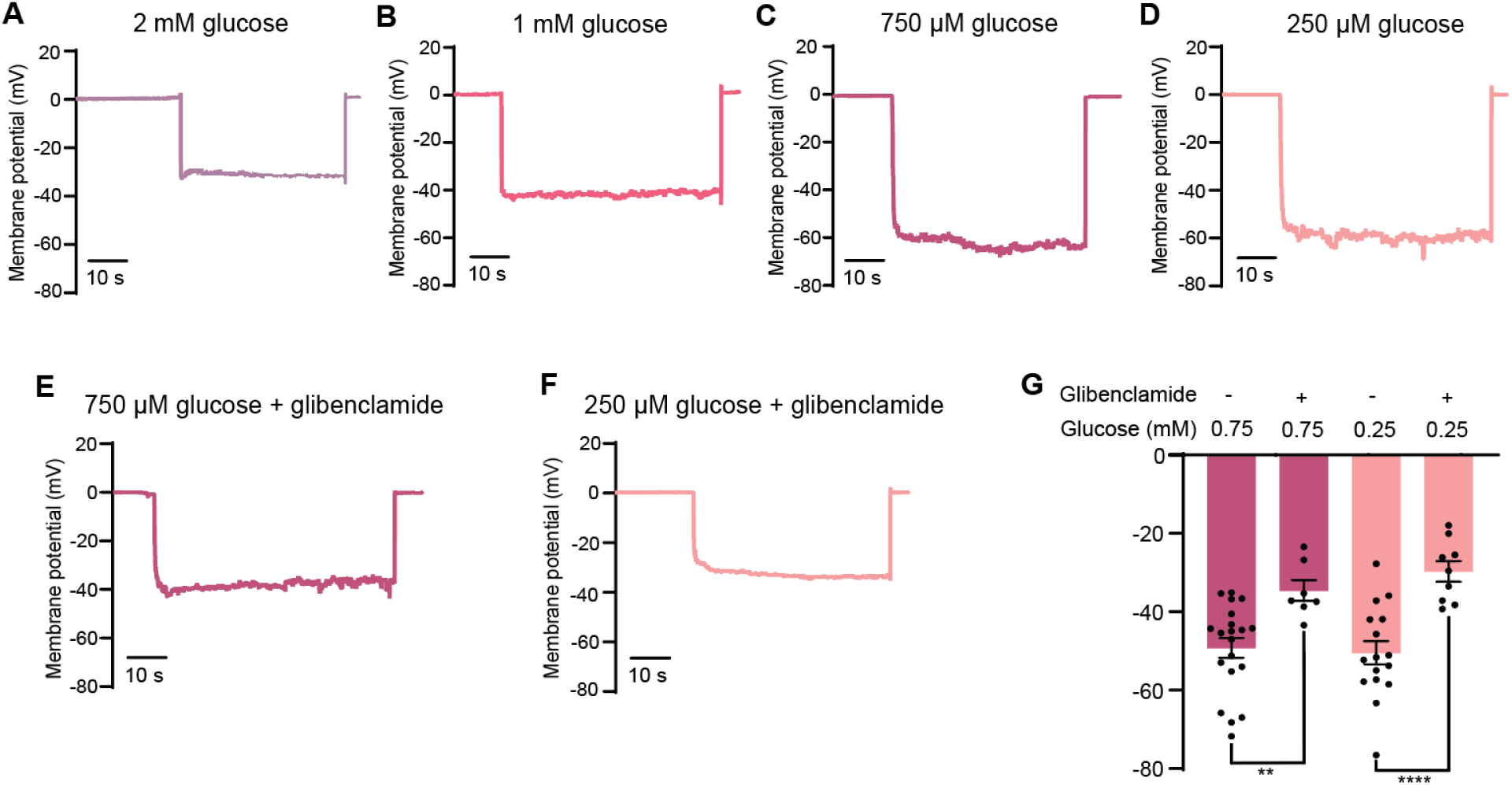
Example traces of Vm measurements with 2 mM bath glucose **(A)**, 1 mM bath glucose **(B)**, 750 μM bath glucose **(C)**, 250 μM bath glucose **(D)**, 750 μM bath glucose with 10 μM glibenclamide **(E)**, and 250 μM bath glucose in the presence of 10 μM glibenclamide **(F). (G)** Summary data showing glibenclamide blocks hyperpolarizing effects of 750 μM and 250 μM glucose (750 μM glucose (20 cells, 4 mice) vs. 750 μM glucose + 10 μM glibenclamide (7 cells, 4 mice): \*\**P* = 0.0094, t_97_ = 3.343; 250 μM glucose (16 cells, 4 mice) vs. 250 μM glucose + 10 μM glibenclamide (9 cells, 4 mice): \*\*\*\**P* < 0.0001, t_97_ = 5.009; One-way ANOVA with Sidak’s multiple comparison test).

**Supplementary movie 1**. Movie depicting imaging area for an experiment in which a deep capillary pericyte was targeted by pressure-ejection of 10 μM pinacidil 170 μm downstream of the imaging site focused on a PA with pre-capillary sphincter and 1^st^ order capillary. Within seconds of remote application of pinacidil, the PA, sphincter and 1^st^ order capillary robustly dilate. Scale bars on Z projections and single-plane recording are 50 and 5 μm, respectively.

## REFERENCES

1. Blinder P, Tsai PS, Kaufhold JP, Knutsen PM, Suhl H, Kleinfeld D. The cortical angiome: an interconnected vascular network with noncolumnar patterns of blood flow. Nat Neurosci. 2013;16:889–897. doi:10.1038/nn.3426

2. Grubb S, Cai C, Hald BO, et al. Precapillary sphincters maintain perfusion in the cerebral cortex. Nat Commun. 2020;11:395. doi:10.1038/s41467-020-14330-z

3. Ratelade J, Klug NR, Lombardi D, et al. Reducing Hypermuscularization of the Transitional Segment between Arterioles and Capillaries Protects Against Spontaneous Intracerebral Hemorrhage. Circulation. 2020;141:2078–2094. doi:10.1161/CIRCULATIONAHA.119.040963

4. Grant RI, Hartmann DA, Underly RG, Berthiaume A-A, Bhat NR, Shih AY. Organizational hierarchy and structural diversity of microvascular pericytes in adult mouse cortex. J Cereb Blood Flow Metab. 2019;39(3):411–425. doi:10.1177/0271678X17732229

5. Gonzales AL, Klug NR, Moshkforoush A, et al. Contractile pericytes determine the direction of blood flow at capillary junctions. Proc Natl Acad Sci U S A. 2020. doi:10.1073/pnas.1922755117

6. Hartmann DA, Berthiaume AA, Grant RI, et al. Brain capillary pericytes exert a substantial but slow influence on blood flow. Nat Neurosci. 2021;24(5):633–645. doi:10.1038/s41593-020-00793-2

7. Hartmann DA, Underly RG, Grant RI, Watson AN, Lindner V, Shih AY. Pericyte structure and distribution in the cerebral cortex revealed by high-resolution imaging of transgenic mice. Neurophotonics. 2015;2(4):041402. doi:10.1117/1.NPh.2.4.041402

8. Berthiaume AA, Grant RI, McDowell KP, et al. Dynamic Remodeling of Pericytes In Vivo Maintains Capillary Coverage in the Adult Mouse Brain. Cell Rep. 2018;22(1):8–16. doi:10.1016/j.celrep.2017.12.016

9. Armulik A, Abramsson A, Betsholtz C. Endothelial/pericyte interactions. Circ Res. 2005;97:512–523. doi:10.1161/01.RES.0000182903.16652.d7

10. Kovacs-Oller T, Ivanova E, Bianchimano P, Sagdullaev BT. The pericyte connectome: spatial precision of neurovascular coupling is driven by selective connectivity maps of pericytes and endothelial cells and is disrupted in diabetes. Cell Discov. 2020;6:39. doi:10.1038/s41421-020-0180-0

11. Ornelas S, Berthiaume AA, Bonney SK, et al. Three-dimensional ultrastructure of the brain pericyte-endothelial interface. J Cereb Blood Flow Metab. 2021:1–16. doi:10.1177/0271678X211012836

12. Armulik A, Genové G, Mäe M, et al. Pericytes regulate the blood-brain barrier. Nature. 2010;468(7323):557–561. doi:10.1038/nature09522

13. Mäe MA, He L, Nordling S, et al. Single-Cell Analysis of Blood-Brain Barrier Response to Pericyte Loss. Circ Res. 2021:E46–E62. doi:10.1161/CIRCRESAHA.120.317473

14. Peppiatt CM, Howarth C, Mobbs P, Attwell D. Bidirectional control of CNS capillary diameter by pericytes. Nature. 2006;443:700–704. doi:10.1038/nature05193

15. Hall CN, Reynell C, Gesslein B, et al. Capillary pericytes regulate cerebral blood flow in health and disease. Nature. 2014;508:55–60. doi:10.1038/nature13165

16. Rungta RL, Chaigneau E, Osmanski B-F, Charpa S. Vascular Compartmentalization of Functional Hyperemia from the Synapse to the Pia. Neuron. 2018;99(2):362–37. doi:10.1016/j.neuron.2018.06.012

17. Zambach SA, Cai C, Helms HCC, et al. Precapillary sphincters and pericytes at first-order capillaries as key regulators for brain capillary perfusion. Proc Natl Acad Sci U S A. 2021;118(26). doi:10.1073/pnas.2023749118

18. Longden TA, Mughal A, Hennig GW, et al. Local IP(3) receptor-mediated Ca(2+) signals compound to direct blood flow in brain capillaries. Sci Adv. 2021;7(30). doi:10.1126/sciadv.abh0101

19. Hariharan A, Weir N, Robertson C, He L, Betsholtz C, Longden TA. The Ion Channel and GPCR Toolkit of Brain Capillary Pericytes (In Press). Front Cell Neurosci. 2020;14. doi:10.3389/fncel.2020.601324

20. He L, Vanlandewijck M, Mäe M, et al. Single cell RNAseq of mouse brain and lung vascular and vessel-associated cell types. Sci Data. 2018;5:180160. doi:10.1038/sdata.2018.160

21. Vanlandewijck M, He L, Mäe MA, et al. A molecular atlas of cell types and zonation in the brain vasculature. Nature. 2018;554(7693):475–480. doi:10.1038/nature25739

22. Tarasov A, Dusonchet J, Ashcroft F. Metabolic regulation of the pancreatic beta-cell ATP-sensitive K+ channel: a pas de deux. Diabetes. 2004;53(Suppl 3):S113–S122. doi:10.2337/diabetes.53.suppl_3.S113

23. Zhao G, Joca HC, Nelson MT, Lederer WJ. ATP-And voltage-dependent electro-metabolic signaling regulates blood flow in heart. Proc Natl Acad Sci U S A. 2020;117(13):7461–7470. doi:10.1073/pnas.1922095117

24. Davis MJ, Hill MA. Signaling mechanisms underlying the vascular myogenic response. Physiol Rev. 1999;79(2):387–423. doi:10.1152/physrev.1999.79.2.387

25. Sancho M, Klug NR, Mughal A, Heppner TJ, Hill-Eubanks D, Nelson MT. Electro-Metabolic Sensing Through Capillary ATP-Sensitive K+ Channels and Adenosine to Control Cerebral Blood Flow. bioRxiv. 2021:2021.03.12.435152. doi:10.1101/2021.03.12.435152

26. Dabertrand F, Nelson MT, Brayden JE. Acidosis dilates brain parenchymal arterioles by conversion of calcium waves to sparks to activate BK channels. Circ Res. 2012;110(2):285–294. doi:10.1161/CIRCRESAHA.111.258145

27. Díaz-Flores L, Gutiérrez R, Madrid JF, et al. Pericytes. Morphofunction, interactions and pathology in a quiescent and activated mesenchymal cell niche. Histol Histopathol. 2009;24(7):909–969. doi:10.14670/HH-24.909

28. Longden TA, Dabertrand F, Koide M, et al. Capillary K+-sensing initiates retrograde hyperpolarization to locally increase cerebral blood flow. Nat Neurosci. 2017;20:717–726. doi:10.1038/nn.4533

29. Tong XY, Porter LM, Liu GX, et al. Consequences of cardiac myocyte-specific ablation of KATP channels in transgenic mice expressing dominant negative Kir6 subunits. Am J Physiol - Hear Circ Physiol. 2006;291(2). doi:10.1152/ajpheart.00051.2006

30. Malester B, Tong X, Ghiu I, et al. Transgenic expression of a dominant negative K ATP channel subunit in the mouse endothelium: effects on coronary flow and endothelin-1 secretion. FASEB J. 2007;21(9):2162–2172. doi:10.1096/fj.06-7821com

31. Damisah EC, Hill RA, Tong L, Murray KN, Grutzendler J. A fluoro-Nissl dye identifies pericytes as distinct vascular mural cells during in vivo brain imaging. Nat Neurosci. 2017;20(7):1023–1032. doi:10.1038/nn.4564

32. Akrouh A, Halcomb SE, Nichols CG, Sala-Rabanal M. Molecular biology of KATP channels and implications for health and disease. IUBMB Life. 2009;61(10):971–978. doi:10.1002/iub.246

33. Brown PD, Davies SL, Speake T, Millar ID. Molecular mechanisms of cerebrospinal fluid production. Neuroscience. 2004;129(4):955–968. doi:10.1016/j.neuroscience.2004.07.003

34. Gruetter R, Novotny EJ, Boulware SD, et al. Direct measurement of brain glucose concentrations in humans by 13C NMR spectroscopy. Proc Natl Acad Sci U S A. 1992;89(3):1109–1112. doi:10.1073/pnas.89.3.1109

35. Choi IY, Lee SP, Kim SG, Gruetter R. In vivo measurements of brain glucose transport using the reversible michaelis-menten model and simultaneous measurements of cerebral blood flow changes during hypoglycemia. J Cereb Blood Flow Metab. 2001;21(6):653–663. doi:10.1097/00004647-200106000-00003

36. Ritter S. Monitoring and Maintenance of Brain Glucose Supply. Appet Food Intake. 2017:177–204. doi:10.1201/9781315120171-9

37. De Vries MG, Arseneau LM, Lawson ME, Beverly JL. Extracellular Glucose in Rat Ventromedial Hypothalamus during Acute and Recurrent Hypoglycemia. Diabetes. 2003;52(11):2767–2773. doi:10.2337/diabetes.52.11.2767

38. Silver IA, Erecinska M. Extracellular glucose concentration in mammalian brain: Continuous monitoring of changes during increased neuronal activity and upon limitation in oxygen supply in normo-, hypo-, and hyperglycemic animals. J Neurosci. 1994;14(8):5068–5076. doi:10.1523/jneurosci.14-08-05068.1994

39. McNay EC, Sherwin RS. From artificial cerebro-spinal fluid (aCSF) to artificial extracellular fluid (aECF): Microdialysis perfusate composition effects on in vivo brain ECF glucose measurements. J Neurosci Methods. 2004;132(1):35–43. doi:10.1016/j.jneumeth.2003.08.014

40. McNay EC, Gold PE. Extracellular Glucose Concentrations in the Rat Hippocampus Measured byZero-Net-Flux. J Neurochem. 1999;72(2):785–790. doi:10.1046/j.1471-4159.1999.720785.x

41. Dunn-Meynell AA, Sanders NM, Compton D, et al. Relationship among brain and blood glucose levels and spontaneous and glucoprivic feeding. J Neurosci. 2009;29(21):7015–7022. doi:10.1523/JNEUROSCI.0334-09.2009

42. Abi-Saab WM, Maggs DG, Jones T, et al. Striking differences in glucose and lactate levels between brain extracellular fluid and plasma in conscious human subjects: Effects of hyperglycemia and hypoglycemia. J Cereb Blood Flow Metab. 2002;22(3):271–279. doi:10.1097/00004647-200203000-00004

43. Longden TA, Hill-Eubanks DC, Nelson MT. Ion channel networks in the control of cerebral blood flow. J Cereb Blood Flow Metab. 2016;36(3):492–512. doi:10.1177/0271678X15616138

44. Patching SG. Glucose Transporters at the Blood-Brain Barrier: Function, Regulation and Gateways for Drug Delivery. Mol Neurobiol. 2017;54(2):1046–1077.

45. Matsunami T, Suzuki T, Hisa Y, Takata K, Takamatsu T, Oyamada M. Gap junctions mediate glucose transport between GLUT1-positive and -negative cells in the spiral limbus of the rat cochlea. Cell Commun Adhes. 2006;13(1-2):93–102. doi:10.1080/15419060600631805

46. Rouach N, Koulakoff A, Abudara V, Willecke K, Giaume C. Astroglial metabolic networks sustain hippocampal synaptic transmission. Science (80-). 2008;322(5907):1551–1555. doi:10.1126/science.1164022

47. Longden TA, Nelson MT. Vascular inward rectifier K+ channels as external K+ sensors in the control of cerebral blood flow. Microcirculation. 2015;22(3):183–196. doi:10.1111/micc.12190

48. Quignard J, Harley E, Duhault J, Vanhoutte P, Félétou M. K+ Channels in Cultured Bovine Retinal Pericytes: Effects of β-Adrenergic Stimulation. J Cardiovasc Pharmacol. 2003;42(3):379–388.

49. Ikematsu N, Dallas ML, Ross FA, et al. Phosphorylation of the voltage-gated potassium channel Kv2.1 by AMP-activated protein kinase regulates membrane excitability. Proc Natl Acad Sci U S A. 2011;108(44):18132–18137. doi:10.1073/pnas.1106201108

50. Fox PT, Raichle ME, Mintun MA, Dence C. Nonoxidative glucose consumption during focal physiologic neural activity. Science. 1988;241:462–464.

51. Lundgaard I, Li B, Xie L, et al. Direct neuronal glucose uptake heralds activity-dependent increases in cerebral metabolism. Nat Commun. 2015;6. doi:10.1038/ncomms7807

52. Hosford PS, Christie IN, Niranjan A, et al. A critical role for the ATP-sensitive potassium channel subunit KIR6.1 in the control of cerebral blood flow. J Cereb Blood Flow Metab. 2019;39(10):2089–2095. doi:10.1177/0271678X18780602

53. McManus R, Ioussoufovitch S, Froats E, St Lawrence K, Van Uum S, Diop M. Dynamic response of cerebral blood flow to insulin-induced hypoglycemia. Sci Rep. 2020;10(1). doi:10.1038/s41598-020-77626-6

54. Fernández-Klett F, Offenhauser N, Dirnagl U, Priller J, Lindauer U. Pericytes in capillaries are contractile in vivo, but arterioles mediate functional hyperemia in the mouse brain. Proc Natl Acad Sci U S A. 2010;107(51):22290–22295. doi:10.1073/pnas.1011321108

55. Alarcon-Martinez L, Villafranca-Baughman D, Quintero H, et al. Interpericyte tunnelling nanotubes regulate neurovascular coupling. Nature. 2020;585(7823):91–95. doi:10.1038/s41586-020-2589-x

56. Payne LB, Hoque M, Houk C, Darden J, Chappell JC. Pericytes in Vascular Development. Curr Tissue Microenviron Reports. 2020;1(3):143–154. doi:10.1007/s43152-020-00014-9

57. Benjamin LE, Hemo I, Keshet E. A plasticity window for blood vessel remodelling is defined by pericyte coverage of the preformed endothelial network and is regulated by PDGF-B and VEGF. Development. 1998;125(9):1591–1598. doi:10.1242/dev.125.9.1591

58. Kisler K, Nelson AR, Rege S V, et al. Pericyte degeneration leads to neurovascular uncoupling and limits oxygen supply to brain. Nat Neurosci. 2017;20:406–41. doi:10.1038/nn.4489

59. Hirunpattarasilp C, Attwell D, Freitas F. The role of pericytes in brain disorders: from the periphery to the brain. J Neurochem. 2019;150(6):648–665. doi:10.1111/jnc.14725

60. Kisler K, Nelson AR, Montagne A, Zlokovic B V. Cerebral blood flow regulation and neurovascular dysfunction in Alzheimer disease. Nat Rev Neurosci. 2017;18(7):419–434. doi:10.1038/nrn.2017.48

61. Girouard H, Iadecola C. Neurovascular coupling in the normal brain and in hypertension, stroke, and Alzheimer disease. J Appl Physiol. 2006;100(1):328–335. doi:10.1152/japplphysiol.00966.2005

62. De La Torre JC. Critically attained threshold of cerebral hypoperfusion: The CATCH hypothesis of Alzheimer’s pathogenesis. Neurobiol Aging. 2000;21(2):331–342. doi:10.1016/S0197-4580(00)00111-1

63. Yang AC, Vest RT, Kern F, et al. A human brain vascular atlas reveals diverse mediators of Alzheimer’s risk. Nature. 2022. doi:10.1038/s41586-021-04369-3

64. Mughal A, Harraz OF, Gonzales AL, Hill-Eubanks D, Nelson MT. PIP2 Improves Cerebral Blood Flow in a Mouse Model of Alzheimer’s Disease. Function. 2021;2(2). doi:10.1093/function/zqab010

65. Jespersen SN, østergaard L. The roles of cerebral blood flow, capillary transit time heterogeneity, and oxygen tension in brain oxygenation and metabolism. J Cereb Blood Flow Metab. 2012;32(2):264–277. doi:10.1038/jcbfm.2011.153

66. Hall CN, Klein-Flügge MC, Howarth C, Attwell D. Oxidative phosphorylation, not glycolysis, powers presynaptic and postsynaptic mechanisms underlying brain information processing. J Neurosci. 2012;32(26):8940–8951. doi:10.1523/JNEUROSCI.0026-12.2012

67. Wei HS, Kang H, Rasheed IYD, et al. Erythrocytes Are Oxygen-Sensing Regulators of the Cerebral Microcirculation. Neuron. 2016;91(4):851–862. doi:10.1016/j.neuron.2016.07.016

68. Erdener ŞE, Tang J, Sajjadi A, et al. Spatio-temporal dynamics of cerebral capillary segments with stalling red blood cells. J Cereb Blood Flow Metab. 2019;39(5):886–900. doi:10.1177/0271678X17743877

69. Matsugi T, Chen Q, Anderson DR. Suppression of CO2-induced relaxation of bovine retinal pericytes by angiotensin II. Investig Ophthalmol Vis Sci. 1997;38(3):652–657.

70. Dienel GA. Brain glucose metabolism: Integration of energetics with function. Physiol Rev. 2019;99(1):949–1045. doi:10.1152/physrev.00062.2017

71. Lee H-W, Xu Y, Zhu X, et al. Endothelium-derived lactate is required for pericyte function and blood–brain barrier maintenance. EMBO J. 2022;e109890. doi:10.15252/embj.2021109890

72. Mosconi L, Pupi A, De Leon MJ. Brain glucose hypometabolism and oxidative stress in preclinical Alzheimer’s disease. Ann N Y Acad Sci. 2008;1147:180–195. doi:10.1196/annals.1427.007

73. Daulatzai MA. Cerebral hypoperfusion and glucose hypometabolism: Key pathophysiological modulators promote neurodegeneration, cognitive impairment, and Alzheimer’s disease. J Neurosci Res. 2017;95(4):943–972. doi:10.1002/jnr.23777

74. An Y, Varma VR, Varma S, et al. Evidence for brain glucose dysregulation in Alzheimer’s disease. Alzheimer’s Dement. 2018;14(3):318–329. doi:10.1016/j.jalz.2017.09.011

75. Xu J, Begley P, Church SJ, et al. Elevation of brain glucose and polyol-pathway intermediates with accompanying brain-copper deficiency in patients with Alzheimer’s disease: metabolic basis for dementia. Sci Rep. 2016;6:27524. doi:10.1038/srep27524

76. Cazzulino AS, Martinez R, Tomm NK, Denny CA. Improved specificity of hippocampal memory trace labeling. Hippocampus. 2016;26(6):752–762. doi:10.1002/hipo.22556

77. Muntifering M, Castranova D, Gibson GA, Meyer E, Kofron M, Watson AM. Clearing for Deep Tissue Imaging. Curr Protoc Cytom. 2018;86(1). doi:10.1002/cpcy.38

78. Susaki EA, Tainaka K, Perrin D, et al. Whole-brain imaging with single-cell resolution using chemical cocktails and computational analysis. Cell. 2014;157(3):726–739. doi:10.1016/j.cell.2014.03.042

79. Longair MH, Baker DA, Armstrong JD. Simple neurite tracer: Open source software for reconstruction, visualization and analysis of neuronal processes. Bioinformatics. 2011;27(17):2453–2454. doi:10.1093/bioinformatics/btr390

